# A model of Ponto-Geniculo-Occipital waves supports bidirectional control of cortical plasticity across sleep-stages

**DOI:** 10.1101/2021.03.16.432817

**Authors:** Kaidi Shao, Juan F. Ramirez Villegas, Nikos K. Logothetis, Michel Besserve

## Abstract

During sleep, cortical network connectivity likely undergoes both synaptic potentiation and depression through system consolidation and homeostatic processes. However, how these modifications are coordinated across sleep stages remains largely unknown. Candidate mechanisms are Ponto-Geniculo-Occipital (PGO) waves, propagating across several structures during Rapid Eye Movement (REM) sleep and the transitional stage from non-REM sleep to REM sleep (pre-REM), and exhibiting sleep stage-specific dynamic patterns. To understand their impact on cortical plasticity, we built an acetylcholine-modulated neural mass model of PGO wave propagation through pons, thalamus and cortex, reproducing a broad range of electrophysiological characteristics across sleep stages. Using a population model of Spike-Time-Dependent Plasticity, we show that cortical circuits undergo different transient regimes depending on the sleep stage, with different impacts on plasticity. Specifically, PGO-induced recurrent cortical activities lead to the potentiation of cortico-cortical synapses during pre-REM, and to their depression during REM sleep. Overall, these results shed light on how the dynamical properties of sleep events propagating to cortical circuits can favor different types of local plastic changes. The variety of events occurring across sleep stages may thus be instrumental in controlling the reorganization of cortical networks from one day to the next.

**Significance statement:** Considerable evidence supports rescaling of cortical synaptic connections during sleep, requiring both long term potentiation to consolidate newly acquired memories, and long-term depression to maintain homeostatic levels of brain activity. However, which aspects of sleep activity contribute to this bidirectional control of plasticity remains unclear. This computational modeling study suggests that widespread transient phenomena called Ponto-geniculo-occipital (PGO) waves, have a sleep-stage dependent effect on plasticity. The alternation between sleep stages can thus be exploited in combination with spontaneously occurring transients, to trigger both up- and down-regulating effects on cortical connectivity, and may explain why the basic structure of sleep-cycles is a well-preserved property across mammalian species.

## Introduction

In most mammalian species, the brain undergoes drastic state changes between different sleep stages – Rapid-Eye-Movement (REM) and non-REM sleep – with broad behavioral and functional similarities. There is moreover an emerging consensus on the existence of sleep-induced plastic changes in relation to two hypothesized functions: memory consolidation, and synaptic homeostasis (Buzsáky, 1998; Tononi and Cirelli, 2003, 2014). However, the detailed underlying mechanisms remain elusive and their investigation requires assessing the brain-wide impacts on plasticity of a variety of phenomena happening during sleep. So far, an extensive amount of experimental and computational modeling studies have focused on the neural bases of transient events observed in non-REM sleep (hippocampal ripples, cortical slow waves and thalamic spindles). In particular, it has been suggested that these non-REM events play a role in long term plastic changes necessary to memory consolidation (Steriade et al., 1993a; Sejnowski and Destexhe, 2000; Puentes-Mestril et al., 2019; Klinzing et al., 2019) and synaptic downscaling (Tononi and Cirelli, 2014, 2020; Watson et al., 2016). However, evidence suggests that key modifications of plasticity not only occur during non-REM but also during REM sleep (Grosmark et al., 2012; Datta and O’Malley, 2013; McDevitt et al., 2015; Boyce et al., 2016, 2017; Zhou et al., 2020).

Interestingly, the transitional stage from non-REM sleep to REM sleep (referred to as pre-REM stage), and the subsequent REM sleep stage are associated with the occurrence of another family of phasic events, Ponto-Geniculo-Occipital (PGO) waves (Steriade et al., 1989; Gott et al., 2017). A typical PGO wave displays biphasic waveforms in Local Field Potential (LFP) activity traces, comprising of a fast-negative component preceding a slower, weaker positive component, possibly followed by smaller fluctuations (Gott et al., 2017; Callaway et al., 1987; Datta, 1997). PGO waves are named after three key structures where their propagation is most frequently observed: the Pons, the Lateral Geniculate Body of the thalamus and the Occipital cortex (Jouvet, 1959). However they also manifest themselves in a broad range of brain structures and a variety of species, e.g. cats (Jouvet, 1959; Hobson, 1965; Datta, 1997), rodents (Kaufman and Morrison, 1981; Datta et al., 1998; Deboer et al., 1998, 1999), non-human primates (Cohen and Feldman, 1968; Ramirez-Villegas et al., 2020) and humans (Lim et al., 2007; Fernández-Mendoza et al., 2009). Importantly, during pre-REM and REM stages PGO waves exhibit two distinct subtypes (Bizzi, 1966): during pre-REM sleep they occur in high-amplitude singlets (Bowker, 1985), whereas REM PGO waves tend to cluster in 3-5 successive weaker deflections (Brooks, 1968; Bowker, 1985; Morrison and Pompeiano, 1966; Datta et al., 1992).

Given that converging evidence supports different roles played by non-REM and REM sleep stages in reorganizing networks across the brain (Grosmark et al., 2012; Seibt et al., 2017), the fact that PGO waves span both sleep stages in the form of two subtypes suggests that these events play a key role in coordinating plastic changes, and their analysis may provide insights into the differences between plasticity promoting mechanisms during non-REM and REM sleep. Indeed, experimental evidence supports a key role of REM-PGO waves in enhancing sleep-dependent learning and memory (Datta, 2000; Datta et al., 2003; Mavanji and Datta, 2003; Datta et al., 2004). In contrast, little is known about the impact of pre-REM PGO waves on plasticity. However, recent electrophysiological evidence for a coupling between hippocampal sharp wave-ripples and PGO waves suggests PGO waves are involved in memory consolidation processes happening during non-REM sleep (Ramirez-Villegas et al., 2020).

In-vivo investigation of PGO-triggered plasticity remains challenging, notably due to the spatial sparsity of the spines receiving PGO-associated potentials, and the expected weaker plastic changes induced by spontaneous activity, in contrast to those triggered by controlled sensory or electrical stimulation (Li et al., 2017; Yang et al., 2014; Zhou et al., 2020). As an alternative, investigating PGO waves from the computational modeling perspective may provide insights and guide further experimental studies. In particular, building a model of large-scale PGO waves propagation may help elucidate how the phasic potentials originated in the pontine region influence widespread brain regions during sleep, and possibly control their plasticity.

In this study, we use a multi-structure neural mass model to reproduce prominent features of PGO waves at a system level and shed light on their possible functions. By including cholinergic neuromodulation in our model, we put an emphasis on reproducing PGO-related phenomena across sleep stages, and account for the variability of PGO wave subtypes occurring either during pre-REM or REM sleep. Finally, we investigate the putative influence on PGO waves on cortical plasticity through a mesoscopic model of Spike Time Dependent plasticity (STDP), suggesting PGO waves may trigger opposite effects in REM and pre-REM sleep. We provide insights on such state-dependent differences are achieved, suggesting a general framework to predict the influence of phasic events on plasticity.

## Results

### Ponto-geniculo-occiptial neural mass model

We designed a neural mass model to simulate average rate-coded population activities of several groups of homogeneous neurons (Wilson and Cowan, 1972, 1973; Jansen et al., 1993; Lopes da Silva et al., 1974) in three structures influenced by PGO waves: the pons, the thalamus (more precisely the lateral geniculate nucleus and the thalamic reticular nucleus) and the primary visual cortex.

Neural mass models represent the state of a population of identical neurons by the average firing rates across this population. The basic elements of such model are illustrated on Fig. 1A. The population average membrane potential evolves according to its intrinsic dynamics as well as post-synaptic currents it receives from connected populations, and outputs the population firing rate as a non-linear instantaneous function of the membrane potential using a sigmoidal activation. After multiplication with a synaptic strength coefficient, the output firing rate is convolved with the impulse response of the synapses that link the population to downstream neurons. Although neural mass models constitute a strong simplification with respect to single neuron models, recent work has shown they can reproduce sleep related phasic patterns, such as spindles and slow waves, with a satisfactory degree of realism, by taking into account key intrinsic currents flowing through the cells’ membrane (Schellenberger Costa et al., 2016). We follow this approach to model PGO-wave propagation through pons, thalamus and cortex.

**Figure 1.**
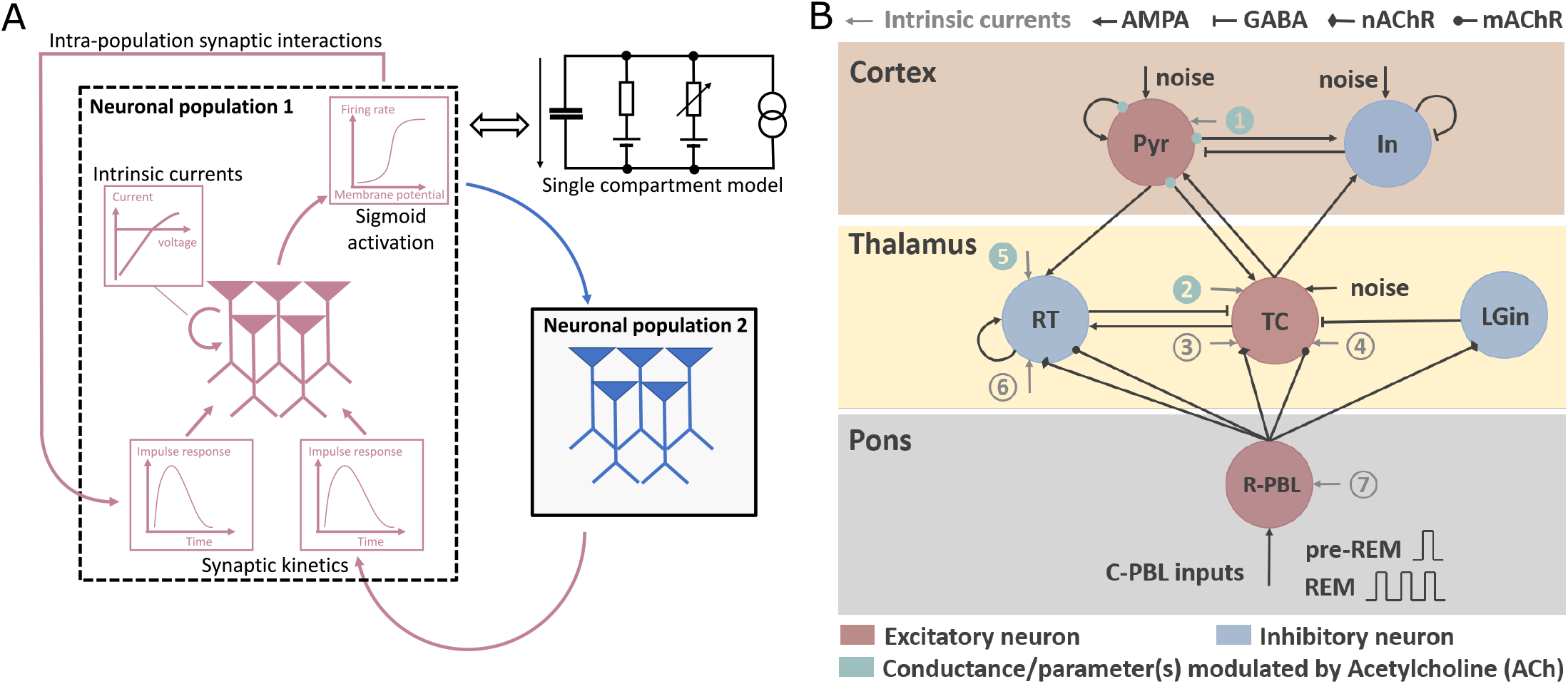
Neural mass model of PGO waves. (A) Illustration of the elements of neural mass models. Two neuronal populations modelled by neural masses are illustrated in separate colors. For a given population (marked in dashed rectangle) intrinsic and synaptic currents drive population membrane potential dynamics, which controls firing rate via a sigmoid activation curve. (B) Global view of the model structure, consisting of biologically plausible interconnected neuronal populations which receive brief pulses as inputs from C-PBL neurons to generate PGO-related neuronal activities. Switch of sleep stages are modulated by 4 major parameters (marked in green) associated with the change of Acetylcholine concentration. The circled numbers indicate intrinsic currents and are listed in Table 1, together with their role in state establishment or PGO wave generation. Abbreviation for neuronal populations: Pyr: Pyramidal neurons; In: inhibitory neurons; TC: thalamocortical neurons; RT: reticular thalamic neurons; LGin: interneurons in LGN; R-PBL: neurons in the rostal peribrachial nucleus (PGO-transferring neurons); C-PBL: neurons in the caudolateral peribrachial nucleus (PGO-triggering neurons). Abbreviation for receptors: AMPA: *α*-amino-3-hydroxy-5-methyl-4-isoxazolepropionic acid; GABA: gamma-Aminobutyric acid; nAChR: nicotinic acetylcholine receptor; mAChR: muscarinic acetylcholine receptor.

Specifically, our model involves 6 neuronal populations, as represented in Fig. 1B. The pontine population (R-PBL), representing low-frequency bursting PGO transferring neurons located in the rostral peribrachial area (R-PBL), receives high-frequency bursts from PGO triggering neurons located in the caudolateral peribrachial area (C-PBL) (Datta, 1997; Datta and Hobson, 1994; Datta, 1995; Datta and Hobson, 1995). The thalamus comprises 3 populations that are hypothesized to underlie PGO wave generation: the thalamocortical relay (TC) neurons in the lateral geniculate nucleus (LGN), the reticular thalamic (RT) neurons in the peri-geniculate nucleus (PGN) (Steriade and Llinás, 1988; McCormick, 1992) and the local intra-geniculate interneurons (LGin) in LGN (Hu et al., 1989a, b). In the cortex, following Schellenberger Costa et al. (2016), we model one excitatory population of pyramidal cells (Pyr), and one inhibitory interneuron population (In).

We model high-frequency bursts in the C-PBL PGO-triggering neurons them as brief pulses — short-lasting rectangular signals — in the population rate activity to reproduce the pooled bursting activities. Based on experimental traces of classical PGO waveforms and firing patterns of C-PBL neurons during both states (Datta and Hobson, 1994), pre-REM PGO waves are appropriately modeled with a single pulse while REM PGO waves are generated by several successive pulses (here we choose 3). The bursting interval (intervals between each two pulses) is fixed and selected from peri-PGO spiking histograms of C-PBL neurons (see also Fig. S1).

We incorporate specific key intrinsic currents in each population in order to model sleep oscilla-tions related to memory consolidation processes, e.g. spindles, slow waves (see Table 1 for a summary and see Materials and Methods for details). In the pons, transferring neurons are equipped with a burst mechanism involving a cholinergic hyperpolarizing current (Leonard and Llinás, 1994). In the thalamus, TC and RT neurons both included a *K*^+^ leaky current and a low-threshold Calcium T-current, together with a hyperpolarization-activated anomalous rectifier h-current in TC neurons (Destexhe et al., 1996; Schellenberger Costa et al., 2016). In the cortex, Pyr neurons possess a sodium-dependent potassium current *I_KNa_* to maintain the non-REM-related cortical oscillations (i.e. slow waves and K-complexes) (Sanchez-Vives and McCormick, 2000).

**Table 1.**
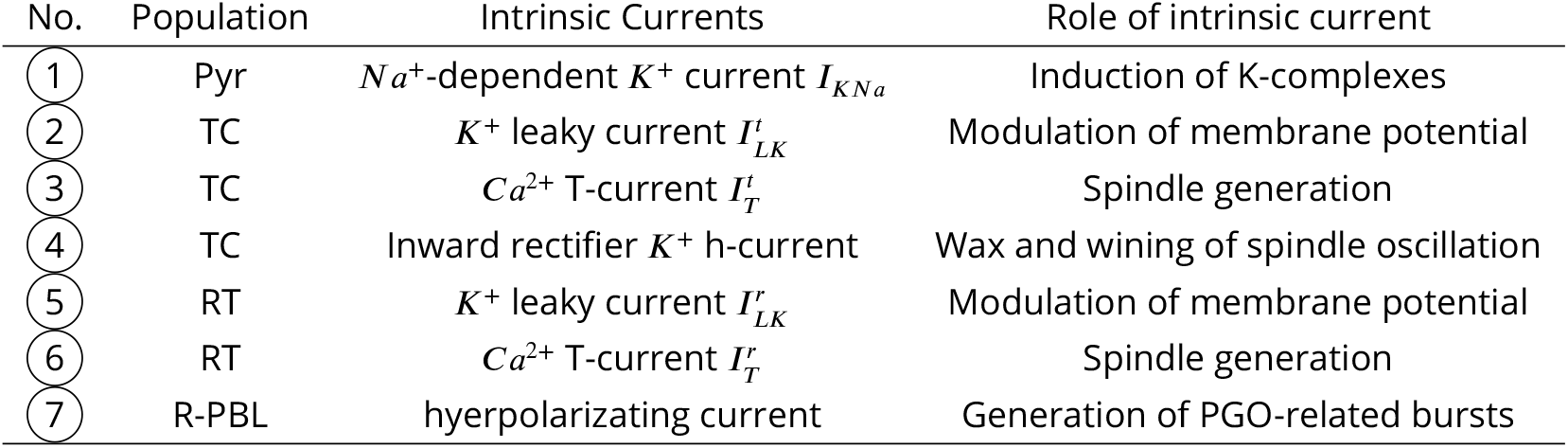
List of intrinsic currents and their corresponding roles in each neuronal population

Neural mass models allow reproducing a wide range of electrophysiological characteristics to compare with the experimental findings. Apart from the population membrane potential and firing rates, we have access to the intrinsic and synaptic currents, enabling us to model the LFPs as a sum of transmembrane currents (Einevoll et al., 2013; Mazzoni et al., 2015) to characterize PGO waveforms. This provides a basis for comparing the outcome of our simulations to PGO waveforms reported in classical studies. Furthermore, following a mesoscopic approximation of the STDP rule (Fung and Robinson, 2013; Robinson, 2011), neural mass models allow making theoretical predictions regarding the plastic changes at a given type of synapse.

### Establishment of non-REM and REM states

In order to model the two PGO wave subtypes appearing in pre-REM and REM sleep, the mechanisms underlying the experimentally observed electrophysiogical changes across sleep stages need to be incorporated in our model. Experimental evidence supports that the transition between non-REM and REM sleep is accompanied by the changing concentration of key neuromodulators, e.g. acetylcholine (ACh) and monoamines (Hobson, 2009; Diekelmann and Born, 2010). From non-REM to REM stages, concentration of the former rises when the latter recedes. Such anticorrelation thus allows us to model transitions between these states only by ACh-dependent influences. More specifically, ACh controls intrinsic conductances mediated by nicotinic/muscarinic acetylcholine receptors (nAChRs/mAChRs) (Hu et al., 1989a; McCormick and Prince, 1986; McCormick and Pape, 1988; McCormick and Bal, 1997). This includes the hyperpolarizing current of R-PBL neurons and potassium leaky currents in both thalamic neurons (see Materials and Methods section Cholinergic modulation of PGO waves for details). In cortex, cholinergic influence is modeled through ACh-dependent parameters modulating population excitability and the sodium-dependent potassium current (Disney et al., 2007; Soma et al., 2012). To match simulated activities to experimental findings, we use the maximum firing rate of the Pyr neurons as the ACh-related critical parameter and tune it to reproduce the peri-PGO firing rates and LFP waveforms of pre-REM and REM stages. Then ACh modulation of each parameter is imposed by a sigmoidal curve achieving a smooth transition between the low-ACh pre-REM value and the high-Ach REM value (see Fig. 2A).

**Figure 2.**
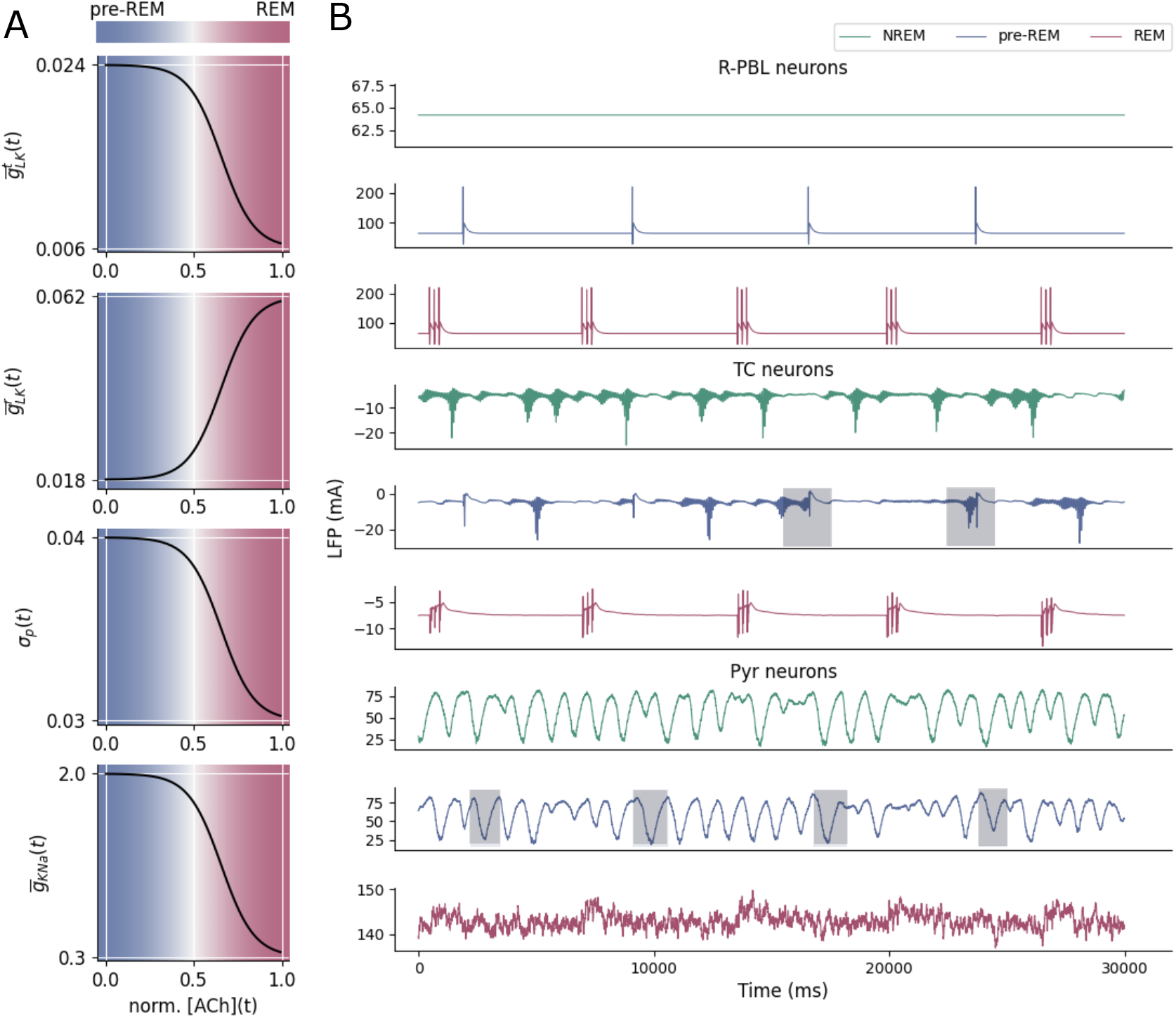
Establishment of sleep stages. (A) ACh-modulated model parameters. 4 critical parameters from the thalamocortical part are selected to reflect the cholinergic influence of the network. Their changes are linked to ACh concentration via a sigmoid-like relationship mimicking smooth stage switches. (B) State comparison of simulated LFPs in R-PBL (top), TC (middle) and Pyr neurons (bottom). LFPs of the three neuronal populations in three states (non-REM, pre-REM and REM) are plotted in separate colors for comparison. During pre-REM state, in the thalamus, contrary to non-REM state, spindles are interrupted by PGOs, as marked by grey shades in the trace of TC neurons during pre-REM stage. In the cortex, PGO waves trigger more slow oscillations during pre-REM, as marked by grey shades in the trace of Pyr neurons in pre-REM stage.

Following this procedure, we first check whether the model reproduces long-term patterns of ponto-thalamo-cortical activity observed experimentally. Fig. 2B shows the LFPs in R-PBL, TC and Pyr neurons simulated in the whole network by tuning the ACh-modulated parameters shown in Fig. 2A, illustrating the contrast between the 3 scenarios: non-REM (pre-REM without PGOs), pre-REM (with PGOs) and REM sleep. During the simulated non-REM stage (Fig. 2B upper panel), R-PBL neurons do not burst, as shown in early studies (McCarley et al., 1978; Sakai and Jouvet, 1980; Steriade et al., 1990). TC neurons show strong spindle oscillations appearing with a frequency of occurrence of 0.1-0.2 Hz in the simulated LFP. One can distinguish two spindle subtypes: type-I spin-dles are accompanied by slow waves in the Pyr neurons reflecting oscillations between two level of membrane potential: UP and DOWN states, while type-II spindles occur in isolation (without cortical slow wave) (Costa et al., 2016). In the presence of PGO inputs (Fig. 2B, top panel), biphasic patterns occur in R-PBL neurons triggered by PGO inputs. In the thalamus, some spindles are blocked by PGO inputs (Fig. 2B, middle panel, shaded areas), matching experimentally-recorded spindle interruption by phasic brainstem stimulations (Hu et al., 1989c). In cortex, pre-REM PGO waves trigger a succession of UP and DOWN states, thereby increasing the amount of slow oscillations with respect to non-REM stage (Fig. 2B, bottom panel, shaded areas). During REM stage (Fig. 2B, lower panel), thalamic spindles disappear with the depolarizing effect of the ACh-modulated current, as well as slow waves in the cortex. The activities of these simulated stages match the corresponding tonic electrophysiological traces shown in Steriade et al. (1989), supporting appropriate modeling of these states.

### Model Reproduction of Pontine and Thalamic neuronal activities

Beyond the sustained activities of pre-REM and REM states, we checked the ability of the model to replicate transient changes in PGO-triggered firing patterns reported by classical literature in cats (Steriade et al., 1989; Datta and Hobson, 1994; Steriade et al., 1990). Fig. S1 shows a comprehensive comparison between the key features of PGO-related neuronal activities in simulations and their experimental counterparts reported in classical electrophysiological studies in both pre-REM and REM stages. In the pons (Fig. S1A), the simulated peri-PGO histograms not only resemble the experimental ones in shape, but temporal characteristics also match quantitatively for bursting duration and onset latency, which are shown in detail in Table S1. See Materials and Methods, section Pontine parameter tuning, for details on the tuning methodology.

In the thalamus, the similarities are reflected in the following aspects (Fig. S1B). First, TC and RT neurons show differences in their baseline firing rates (indirectly membrane potentials), reflecting cholinergic modulation of thalamic membrane potentials via a potassium leaky conductance. TC neurons are more depolarized during REM than during pre-REM, while this is the opposite for RT neurons. As depolarization inactivates the T-curents of TC cells, state modulation also switches the dynamics of TC neurons from bursting during pre-REM, modeled by the sharp increase in firing rate at PGO wave onset, to non-bursting during the simulated REM state. In contrast, consistent with experimental studies (Steriade et al., 1989), RT bursting activity spans both sleep stages. Besides, on a finer scale, after bursting RT neurons undergo a slower hyperpolarization induced by the mAChR-receptor-mediated synapse, which is reproduced by the simulated events. Such hyper-polarization is also present during REM stage, contributing to the decreased response in RT neurons to the second and third pontine pulses. In line with experimental evidence, TC neurons show prolonged increased firing following their initial bursts (McCormick and Prince, 1987a; McCormick, 1992). Such sustained firing patterns are caused by activation of the cholinergic ponto-thalamic synapses mediated by a mAChR receptor, together with the dis-inhibition effect by RT neurons during their hyperpolarization (Hu et al., 1989b). In addition, the similarity between experimental and simulation results is significant (permutation test, p<0.05), as measured by a larger cosine similarity between PGO peri-event time windows in comparison with those from randomly selected trials as shown in Fig. S1D. These similarities all validate the reliability of the model for replicating cellular in-vivo activity, supporting it can serve as a steppingstone to investigate plasticity induced by PGO wave activities.

### Model Reproduction of thalamocortical LFP

We further checked the model is also able to reproduce the typically observed PGO-triggered LFP waveforms in the thalamus and cortex, which were more frequently recorded in early reports of PGO waves. A comparison of the LFP waveforms and corresponding spectrograms for the two subtypes of PGO waves is shown in In Fig. 3A-B. The simulated thalamic activity can capture many features of the experimental waveforms (compared to traces in e.g. Callaway et al. (1987)), such as the biphasic pattern consisting of the negative and positive deflections, as well as the stronger amplitude of the negative peak of the pre-REM subtype. Moreover, the duration of the simulated waveforms (approximately 500ms) matches the experimentally recorded waveforms (e.g. in Callaway et al. (1987); Steriade et al. (1989); Nelson et al. (1983)). Spectrograms of the two subtypes in both thalamus and cortex also show that the pre-REM PGO waves are prominent in lower frequency bands compared to REM PGO waves. The reproduction of these electrophysiological characteristics may depend on the parameter tuning of the five ponto-thalamic projections cooperatively transforming the pulse-like pontine outputs into the subtypes’ specific waveforms in the thalamus. We investigated whether all of these projections play a role in the waveform shapes by comparing our result to simulations resulting from suppressing one projection at a time (see Materials and Methods, section Validation of Ponto-thalamic projections). Overall, this analysis supports that taking into account all ponto-thalamic projections is necessary to replicate the REM-PGO waveforms (see Fig. S2).

**Figure 3.**
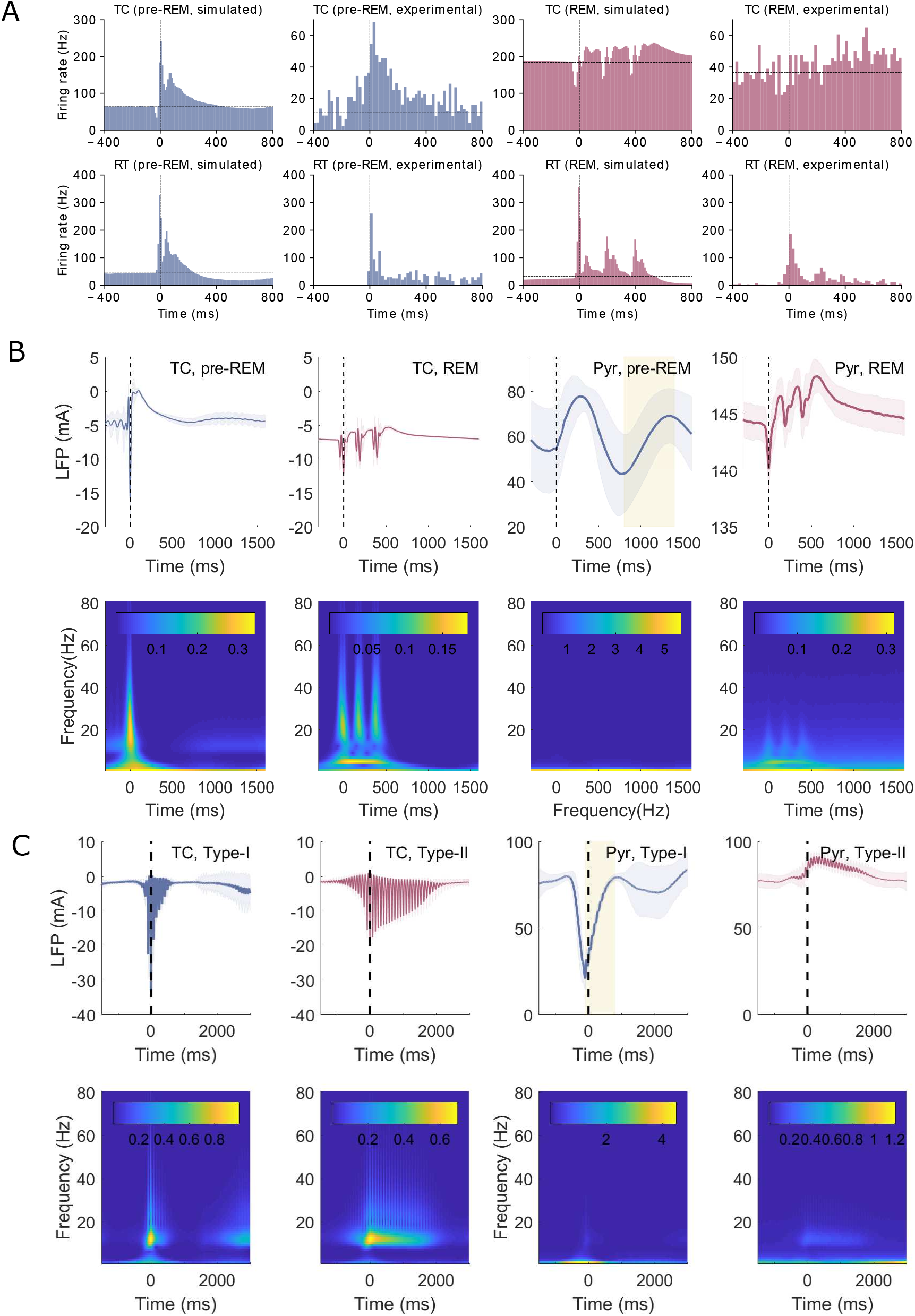
Model Reproduction of thalamocortical LFP. (A) Averaged events and normalized peri-event spectrogram of TC and Pyr neurons during pre-REM and REM PGO waves. Half-transparent shades represent the standard deviation of time-varying events across 1000 trials. Yellow shades mark the DOWN→UP state transition in the cortex. (B) Same as A for type-I and type-II spindles. (C) Comparison of normalized power spectrum for all the conditions. Power spectrum are computed by an average across time and normalized by frequency-wise standard deviation. Shades reflect variability across 1000 trials (trial-wise standard deviation).

We additionally investigated the differences between the PGO waveforms and another key sleep event, observable in thalamus and cortex during NREM sleep, the spindles. Comparing spectrograms and the averaged spectra (Fig. 3B-C), spindles in the thalamus oscillate in a higher frequency band than both PGO waves. Moreover, while Type-I spindles and pre-REM PGO waves are both associated to DOWN-to-UP state transition in the Pyr neurons of the cortex, spindles occur after an UP-to-DOWN transition is initiated by cortex, while pre-REM PGO waves first initiate an UP-state followed by a slow oscillation between DOWN and UP states. This observation, as well as the time scale (200-300ms), matches well with experimental recordings showing that stimulation of the cortex triggers DOWN→UP transitions (Levenstein et al., 2019).

### PGO-triggered cortical plasticity

After this comprehensive validation of the model’s ability to reproduce many electrophysiological aspects, we use the simulated PGO-triggered activities to explore the plasticity effects that PGO waves may induce in the cortical pyramidal neurons. We focus on the plasticity of cortico-cortical excitatory connections, which are expected to be modified by memory consolidation and homeostatic processes during sleep. The considered presynaptic current is thus the excitatory AMPA current *J_Pyr–Pyr_* that the Pyr population sends to itself, and the Pyr post-synaptic firing rate *Q_Pyr_* measures the postsynaptic activities (Fig. 4A).

**Figure 4.**
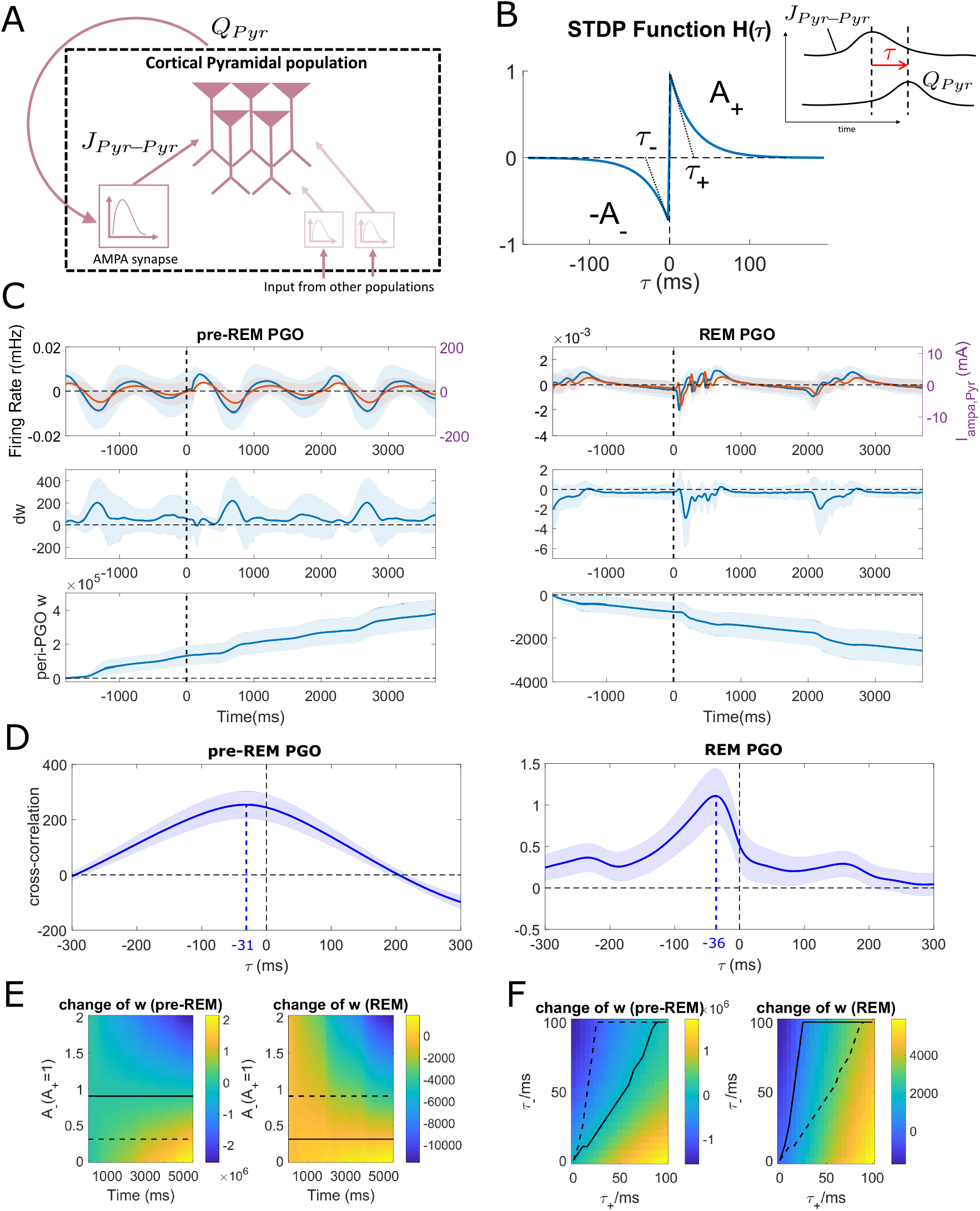
PGO-triggered cortical plasticity. (A) Illustration of the relationship between pyramidal pre-synaptic current *J_Pyr–Pyr_* and post-synaptic firing rate *Q_Pyr_* controlling cortico-cortical STDP plasticity. The impulse response of the AMPA synapse links the two quantities. (B) Illustration of the STDP window. The horizontal axis *τ* represents the time difference between pre- and post-synaptic activities (see right inset). The synapse gets strengthened when pre-synaptic actvities elicit post-synaptic ones, and vice versa for the opposite sequence. (C) Smoothed change of synaptic strength in the intra-cortical excitatory synapse evoked by two subtypes of PGO waves. (top) demeaned waveforms of pre-synaptic current (red) and postsynaptic firing rate (blue). (middle) time-varying change of synaptic strength. (bottom) synaptic strength of the intra-cortical excitatory synapse changing with time. (D) Comparison of cross-correlation between the pre-synaptic current and the post synaptic activities. Blue shades represents standard deviation at each lag across all PGO events (N=1000). (E) Effect of STDP parameter *A*^−^ on plasticity induced by each PGO subtypes. Color bar indicates the relative increase/decrease of synaptic strength across time. Solid lines marks the critical value of the parameter that switches plasticity direction, i.e. potentiation v.s. depression;dashed lines correspond to the critical values of the other subtype. (F) Effect of STDP parameters *τ*_+_ and *τ*^−^ on plasticity induced by the two PGO subtypes. Color bars and lines are analogous to E.

Following a plasticity framework for mean-field models proposed by Robinson (2011) and Fung and Robinson (2013), we compute a time-dependent plasticity rule for neural mass models. The change of synaptic strength *N_Pyr–Pyr_* depends on the simultaneous change of synaptic current *J_Pyr–Pyr_* and postsynaptic firing rate *Q_Pyr_*:

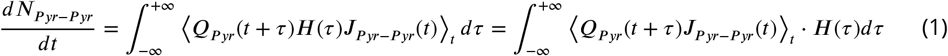

where *H*(*τ*) is the classical STDP function shown in Fig. 4B and the bracket denotes a time averaging. The STDP function’s parameters are taken from in vitro measurements in hippocampal cell cultures (Bi and Poo, 1998). Simply put, this equation shows that the sign and strength of plastic changes are determined by the similarity between the shape of the classical STDP function (Fig. 4B) and the shape of the cross-correlation function between the pre- and post-synaptic activities, measured by the integral of their product overtime. We evaluated the weight changes triggered by the STDP rule in response to the PGO-triggered cortical AMPA current and the firing rate of Pyr neurons based on 1000 simulated PGO waves of each subtype (see Methods, section Computation of triggered plasticity for details). Interestingly, the synaptic modifications differ strongly across PGO waves subtypes. The strength of intra-cortical excitatory synapses rises sharply during pre-REM PGO waves, suggesting that pre-REM PGO waves are important contributors of LTP during this sleep-stage (Fig. 4C, left panel). In contrast, REM-PGO waves elicit Long-term Depression (LTD) in the same synapses (Fig. 4C, right panel).

To interpret the differentiated plastic effects induced by the two PGO subtypes, the cross-correlation function between the pre- and post-synaptic activities exploited by the STDP rule implementation is shown in Fig. 4D. The similar location of the maximum cross-correlation lag(−31ms and −36 for pre-REM and REM PGO waves, respectively), achieving a negative value for both subtypes, reflects that the considered presynaptic AMPA current corresponds to a filtered version of the postsynaptic firing rate through the convolution with a causal filter modeling the synaptic dynamics (see Methods section Modeling methodology). However, another characteristic of the cross-correlation differs between the pre-REM and REM PGO events: the cross-correlation in pre-REM is much flatter than during REM, explaining the sign difference of plasticity. Indeed, the flat pre-REM cross-correlation function implies that equation 1 is approximately proportional to the integral of the STDP function of Fig. 4B. Because this function has a larger positive area (in the positive lags) than the negative, this leads to an overall potentiating effect. For the REM case, the sharp peak of the cross-correlation function at small negative lags puts more weight in the negative portion of the STDP function of Fig. 4B in equation 1, resulting in overall depressing effect on the synapse.

We next investigated the sensitivity of these results to the parameters of the STDP function: the positive amplitude *A*^+^, the negative amplitude *A*^−^, the time constants *τ*_+_ and *τ*_−_. The sign and relative magnitude of plastic changes being unchanged by an overall rescaling of the STDP function, we fix the positive amplitude to a constant *A*^+^ = 1 for this analysis. Fig. 4E-F shows how the switch between LTP and LTD is modulated by changing these parameters. The region comprised between the solid and dashed line shows the domain of STDP function parameters ensuring pre-REM PGO waves are accompanied with an increase of synaptic strength while REM PGO waves occur with a decrease of synaptic strength. These regions are further described in Fig. S3. As demonstrated by our above results, the STDP function parameters measured by (Bi and Poo, 1998) fall into this region (*A*^+^ = 1; *A*^−^ = 0.75, *τ*^+^ = *τ*^−^ = 20*ms*).

In order to interpret the dynamics of other memory-relevant sleep events from a cortical plasticity perspective, we also applied the same STDP framework to the two subtypes of spindles simulated during non-REM stage (Fig. S4). Type-I spindles induce LTP-like behavior in the cortico-cortical connections, which is expected as the peri-spindle activities exhibit a similar pattern as pre-REM PGO waves. Interestingly, type-II spindles lead to a much weaker (see axis scale) temporal increase ofthe synaptic strength before it returns to a lower level. This analysis supports that type-II spindles induce a weaker LTP than type-I. This is in agreement with experimental calcium imaging data indicating that type-I spindles lead to stronger increases of dendritic calcium (prone to trigger large plastic changes) compared to isolated (type II) spindles (Niethard et al., 2018). Our analysis suggests that spontaneous sleep events occurring in the cortico-thalamic system are potentiating the cortico-cortical connections during non-REM sleep (including pre-REM) and depressing during REM sleep, implying a complementary role for these stages in regulating cortical network modifications.

## Discussion

In this study, we have shown that an acetylcholine-modulated ponto-thalamo-cortical neural mass model spanning three sleep stages, non-REM, pre-REM and REM, can reproduce a series of electrophysiological features associated with PGO waves, including the differences in firing rate patterns and LFP waveforms of two PGO wave subtypes. Analysis of cortico-cortical plasticity associated to these events, as well as spindles, suggests a sleep-stage dependent role: non-REM and pre-REM sleep events inducing long-term potentiation, while REM events lead to long-term depression.

The choice of a neural mass model equipped with intrinsic currents helps to maintain the realism of the model by trading off the complexity that would have incurred using individual units. It possesses the ability to account for detailed properties of synaptic and intrinsic currents based on quantitative experimental results, in particular the acetylcholine-modulated state-switching conductance and the five ponto-thalamic cholinergic projections, as the key of the generation of PGO-triggered neural activities. By accounting for the detailed cellular mechanisms of thalamic PGO wave generation, this models contributes to addressing questions debated in the classical electrophysiological literature (Hu et al., 1989a; Datta, 1997). In particular, our model supports the contribution of local geniculate interneurons receiving pontine input to the PGO phenomenon, while they are often neglected in previous thalamocortical models (Muller and Destexhe, 2012). Overall, the insights into the relevant cellular mechanisms provided by this work pave the way for future single unit models of PGO waves. The STDP framework for neural field theory, which has proved effective in characterizing TMS-induced plasticity (Wilson et al., 2014, 2018), is extended here to neural mass models deprived from spatial structure. The STDP rule, computed with parameters estimated experimentally in-vivo (Bi and Poo, 1998), reveals an opposite plasticity effect of two subtypes of PGO waves - potentiation for pre-REM PGO waves and depression for REM PGO waves. These results support the ability of PGO waves to up- and down-scale cortical synaptic weights in larger proportions than the baseline activity surrounding them, making them candidates to enforce synaptic homeostasis, a role that spontaneous phasic sleep events are hypothesized to play in line with considerable experimental evidence (Niethard et al., 2018; Gridchyn et al., 2020). Our results for REM PGO waves are in line with the downscaling effect of REM sleep on cortical spike rates reported in rats (Watson et al., 2016), and the elimination of spines during REM sleep reported in the mouse (Zhou et al., 2020). On the other hand, the potentiating effects that we found for pre-REM PGO waves are in line with converging evidence suggesting non-REM events may be involved in consolidating specific memory traces into local neural microcircuits, with the slow-wave-ripple coupling a specific example (Maingret et al., 2016; Latchoumane et al., 2017; Klinzing et al., 2019). Interestingly, Ramirez-Villegas et al. (2020) reported the co-occurrence of pre-REM PGO waves with sharp-wave ripples, while the present results support a co-occurrence of pre-REM PGO waves and slow waves (specifically DOWN→UP state transitions). According to synaptic plastic pressure theory proposed by Levenstein et al. (2017), ripples, associated with off-line reactivation of episodic memories, occurring during DOWN→UP state transitions provide an opportune time window for new memory traces with low average firing rates to get assimilated into the existing network with relatively high firing rates. Pre-REM PGO waves may serve as trigger and facilitator of this procedure.

Overall, our results suggest the variability of phasic sleep events observed during sleep, may serve the purpose of differentially affecting the plasticity of network elements. The idea that specific phenomena are optimal to trigger certain plasticity mechanisms in cortical circuits resonates with recent experimental results showing spindles induce dendritic depolarization independent of somatic activity, resulting in plasticity promoting calcium activity in cortical dendrites during non-REM sleep (Seibt et al., 2017). To complete our understanding, further experimental work should be conducted to characterize the impact of transient phenomena during REM activity, and must be paralleled with a detailed theoretical investigation of the optimality of transient events for triggering specific plastic changes in cortical circuits. Answering these question holds potential for designing optimal cortical stimulation that maximize the restoration of damaged brain networks, in line with recent evidence that sleep-like oscillatory activities may promote post-stroke recovery (Facchin et al., 2020; Ramanathan et al., 2018).

## Materials and Methods

### Modeling methodology

We provide below the details of our model. The reader can refer to Appendix A: modeling assumptions in SI for further justification based on the literature.

#### Neural mass models

Neural mass models reflect average rate-coded population activities of a group of homogeneous neurons (Jansen et al., 1993; Lopes da Silva et al., 1974), as illustrated in Fig. 1A. Although neural mass models constitute a strong simplification with respect to single neuron models, recent work has shown such model can reproduce sleep related phasic patterns with a satisfactory degree of realism by taking into account intrinsic currents flowing through the cells’ membrane (Schellenberger Costa et al., 2016).

#### Firing rates

The firing of a given population of neurons is assumed to be elicited when its average membrane potential goes beyond a threshold. Thus in a simplified implementation the firing rate *Q_k_* of a neuron population *k* is a sigmoidal function of its instantaneous membrane potential *V_k_*:

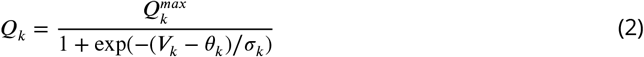

In this function, *θ_k_* denotes the physiological activation threshold which can be obtained from experiments; *σ_k_* is the activation gain that is often influenced by neuromodulators and more generally the brain state.

#### Intrinsic currents

Both intrinsic currents and synaptic currents affect the cells’ membrane potentials. Modeling intrinsic currents amounts to modeling the membrane’s ion channels because they are only driven by the membrane itself instead of being activated by presynaptic neurons. The current that passes through a channel *i* in population *k* is classically expressed as the product of a fixed maximum conductance 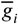 (representing the conductance when the channel is completely open) with a potential difference between the membrane potential *V_k_* and the reversal potential *E_i_* of the channel (as a driving force). Depending on the electrophysiological characteristics of a channel, sometimes an additional factor is used to model a voltage- (or ion-concentration-) dependent opening probability of the channel. As a consequence, a voltage-dependent intrinsic current from channel *i* in a population *k* is modeled in the form:

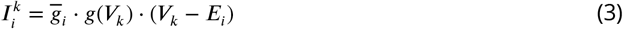

An important intrinsic current that controls the resting membrane potentials of TC and RT neurons is the *K*^+^ leaky current (Krishnan et al., 2016). The voltage-independence of its opening probability is implied by its name, as “leaky” refers to the channels with a linear I-V curve. Thus in population *k* this leaky current (with reversal potential *E_k_* for *K*^+^) is denoted 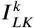 and modeled as:

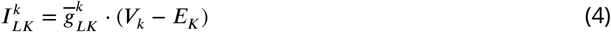

In addition, both TC and RT neurons possess a low-threshold Calcium T-current that generates rebound bursts in their firing patterns. The bursts are generated because the de-inactivation threshold of a T-current is lower than the resting membrane potential - it can only be de-inactivated upon hyperpolarization, so that bursting activities are caused by the overlapping time regime between the voltage-dependent gating of activation and inactivation. The modeling of T-currents in population *k* follows a Hodgkin-Huxley formulation, i.e. the opening probability is written in the form of products of two voltage-dependent gating variables 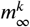 and 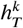 representing respectively the activation and inactivation variables of this Calcium channel (see Destexhe et al. (1998) for more details).

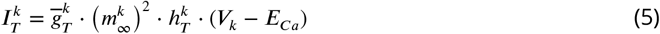

The T-currents in the two neuron types show distinct characteristics in bursting frequency and acceleration. The underlying cellular differences involve (in)activation potentials and time constants of the gating variables, which were already investigated and quantified in earlier studies (Destexhe et al., 1994, 1996, 1998).

The h-current in TC neurons, controlling the waxing and waning of spindles is also called an anomalous inward rectifier because its conductance decreases with rising of membrane potential (Steriade and Llinás, 1988). Following what was implemented in the original model, we assume that the opening probability change is due to extracellular *Ca*^2+^ concentration, whereas the time constants were fitted using experimental results (Destexhe et al., 1996):

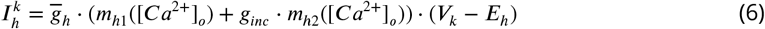

In the cortex, the generation of slowwavesand K-complexes is mediated by a sodium-dependent potassium current in Pyr neurons, where the conductance depends on sodium concentration [*Na*]:

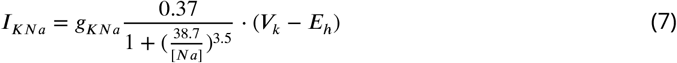

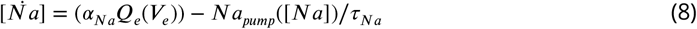

#### Postsynaptic currents

Apart from intrinsic currents, the action potentials of afferent populations of neurons also affect the membrane potential dynamics through a variety of chemical synapses. In the typical case of chemical synapse, neurotransmitters released due to the activation of the presynaptic neuron diffuse in the synaptic cleft and lead to the opening of targeted ion channels on the postsynaptic membranes.

Here we assume a monosynaptic connection *m* from presynaptic neuron population *k′* to postsynaptic population *k*. Then a synaptic current 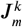 obeys

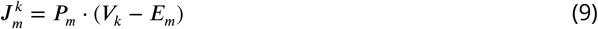

where *P_m_* is the opening probability of postsynaptic channels. Unlike intrinsic currents, *P_m_* is not only dependent of the target population’s own membrane potentials *g*(*V_k_*), but also on the instantaneous concentration of released neurotransmitters, i.e.

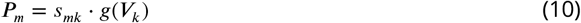

where *s_mk_* represents the opening probability caused by neurotransmitters.

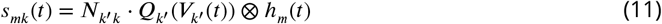

Therefore the complete model for the synapse can be written as

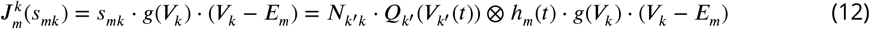

Compared to the general form of intrinsic currents, the product of the synaptic strength and maximum firing rate of presynaptic neuron 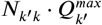 can be seen as equivalent to the maximum conductance (Eq. 2). It is general be free to adjust *N_k′k_* as it depends on many biological parameters of the network for which we do not have a reliable estimate (synaptic strenghts, number of synapses per cell, etc.). The impulse response *h_m_* can be approximated using the time course of postsynaptic currents (PSCs) in response to brief pulses that can be obtained in voltage-clamp experiments. A common approach to approximate such impulse responses is to model them as an alpha function:

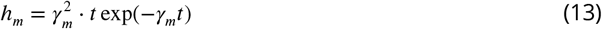

The alpha function reaches its peak value atthe time point 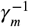. Such impulse response corresponds to a second order linear dynamical system. As a consequence, the opening probability can be written in the form of a differential equation (for derivation, see Appendix. A):

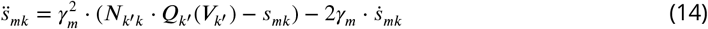

which is thus easy to simulate using classical iterative methods.

Most experiments revealing the PSC of an ion channel characterize its kinetics with a rise time and a decay time, which is imcompatiable with the alpha function. To incorporate these temporal features, an alternative impulse response could be a concatenation of two first-order linear systems with different time constants (referred later as ‘two-exponential’). In the time domain, it take the following form:

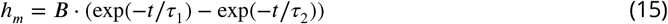

where *B* is the normalization term:

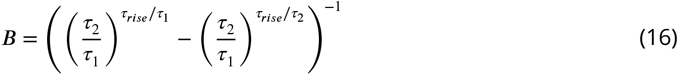

In this formula, *τ_rise_* denotes the rise time, whereas the decay time is set by *τ*_1_ · *τ*_2_ is calculated via the relationship *τ_rise_* = *τ*_1_*τ*_2_/(*τ*_2_ – *τ*_1_). This impulse response reaches its maximum at time ln(*τ*_1_/*τ*_2_) · *τ_rise_*. Similarly to the case of alpha function, the convolution by *h_m_* can be written in the form of a differential equation:

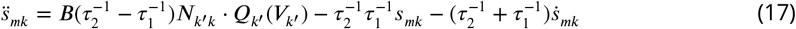

The ‘two-exponential’ assumption of synaptic kinetics captures more characteristics, but induce one more parameter. Therefore, to limit the number of unknown parameters, we model all synapses with precisely-reported time constants with the’two-exponential’framework but confine unknwon synapses within the alpha function framework.

For synapses with long-distance projections across brain regions (e.g. thalamocortical projection), following the original paper, we deal with the axonal conductance delay by adding another linear filter of the synaptic output, which can be approximated by the convolution of another alpha function:

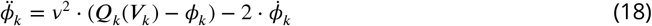

Membrane potential and local field potential (LFP)

The final step is to establish how the membrane potential evolves with the transmembrane currents. The neuron population is assumed equivalent to a neuron modelled with a single compartment model. As illustrated in Fig. 1, the most simplified case is a neuron population *k* influenced by a single (*m*-type) synaptic current 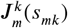 and a single intrinsic current *I_i_*.

The denoted synaptic conductance follows the representation in Eq. 10. With Kirchhoff’s current law, we would be able to get the adaptation of membrane potential *V_k_* with currents:

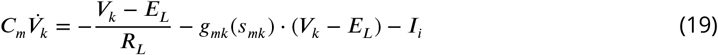

The second term on the right in Eq. 19 represents the synaptic current, where *g_mk_*(*s_mk_*) is the real synaptic conductance. Comparing this form with the synaptic current model (Eq. 9, 10), the synaptic conductance *s_mk_* · *g*(*V_k_*) can be seen as the real conductance *g_mk_*(*s_mk_*) normalized by the leaky conductance 1/*R_L_*, i.e.

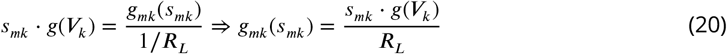

Thus combining Eq. 9, 10, Eq. 19 can be rewritten as:

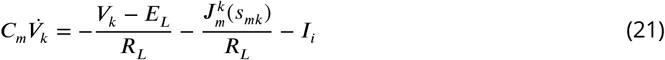

By rearranging the denominator, we can obtain:

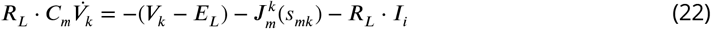

The first term on the right in Eq. 22 can be understood as another equivalent current 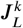 that passes through the membrane. 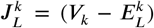 is defined as the general leaky current, a linear current unaffected by either presynaptic or postsynaptic population. Note the *K*^+^ leaky current is excluded because we mentioned in Section Cholinergic modulation of PGO waves that it is influenced by the concentration of ACh.

In experiments a commonly reported feature is the membrane time constant *τ_k_* = *R_L_* · *C_m_*. With a rearrangement of terms, Eq. 22 can be written as:

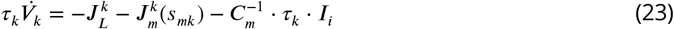

In more general cases, the numbers of synaptic as well as intrinsic currents are not necessarily confined to one. For multiple currents we can obtain the general form:

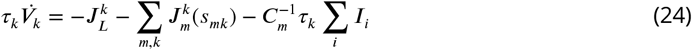

In many experimental studies, the thalamic PGO waves are often characterized by their LFP traces. While the detailed contribution of surrounding transmembrane currents to the LFP is typically unknown, one study attempted to link LFP with proxies based on currents flowing through the surrounding neuronal populations (Mazzoni et al., 2015). In this study the authors approximate the LFP with a weighted sum of synaptic currents, e.g. AMPA and GABA currents, which is to some extent compatible with our model. However, in our model the intrinsic currents also play important roles. We propose to model the LFP of a neuron population as a instantaneous sum of all currents to take into account the effects of intrinsic current:

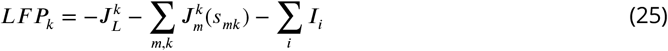

#### Cholinergic modulation of PGO waves

The levels of acetylcholine concentration in the pons and in the thalamus are mainly influenced by the cholinergic tonic firing neurons in the R-PBL which directly project to both thalamic nuclei. From non-REM to REM states, these neurons continuously increase their firing rates (Steriade et al., 1990). We use this observation to build a simple linear approximation of the transition between the two states depending on the normalized ACh concentration [*ACh*](*t*) (ranging from 0 to 1).

Our strategy to link ACh concentration to the activity patterns of PGO-related neurons is as follows:

- First, pick several crucial parameters in the model based on biological plausibility (e.g. maximum conductance for currents or activation threshold of neuronal populations), which were reported to contribute to the switch from non-REM sleep to REM sleep.
- Next, adjust and fix these parameters to reproduce the firing modes of neurons in pre-REM and REM states.
- Finally, establish a sigmoidal relationship between the ACh concentration and the crucial pa-rameters based on experimental evidence, i.e. highest ACh concentration during REM and lowest during pre-REM.

For the cortex, the state-reconstruction can resolved by simplifying the neuromodulated isolated cortical model in (Costa et al., 2016): instead of modulating the cortical network with the concentrations of ACh, serotonin and noradrenaline, we restrict the neuromodulation to ACh. As for the cortical ‘critical parameters’, we followed their choice of the adaptation strength of the sodium-dependent potassium current 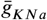, but changed the other to the maximum firing rate of Pyr neurons 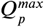 (Barkai and Hasselmo, 1994; Disney et al., 2007; Soma et al., 2012; Steriade et al., 2001) to ensure a broader range of REM activities.

ACh has been reported in vitro and in vivo to tonically depolarize TC neurons (McCormick and Prince, 1987a; Deschênes and Hu, 1990; Steriade et al., 1993b)and hyperpolarize RT neurons, both via a mAChR-mediated *K*^+^ channels (Varela and Sherman, 2007; McCormick and Prince, 1987a; Varela, 2014). State transitions in the thalamus are implemented by modulating the *K*^+^ leaky conductance present in the model of both TC and RT neurons, as described in Section Intrinsic currents.

ForTC neurons, we decreased its potassium leaky conductance conductance from 0.24 to 0.06 to mimick its depolarization. The same conductance in RT neurons was increased from 0.18 to 0.62 to simulate the hyperpolarization. The values of conductance are optimal parameter according empirical experience of tuning the dynamic system, whereas various other combinations would also satisfy the requirements for state switches.

While some related studies (McCormick and Prince, 1987a; Deschênes and Hu, 1990; Zhu and Uhlrich, 1998), measuring how much the leaky conductance was changed in response to ACh application, might have provided a quantitative basis for setting the conductance values, we did not follow them because many unknown experimental variables (in vivo/in vitro, anesthesia, doses of micro-injections, etc.) lead to large uncertainty on the parameter choice.

### Pontine parameter tuning

The quantitative match observed in Fig. S1A results from appropriate adjustment of the conductance of the hyperpolarizing current in R-PBL neurons, which control its bursting behavior. The cellular mechanisms at play and resulting parameter choice are illustrated in Fig. S1C. Bursting activities are co-activated by a low-threshold calcium T-current (Kang and Kitai, 1990; Kamondi et al., 1992; Leonard, 1990) and a hyperpolarizing mAChR-mediated intrinsic cholinergic current (Leonard and Llinás, 1994). The low-threshold T-current, similar to the standard T-current discovered in the thalamus and fitted on rat electrophysiological data, is activated only upon resting hyperpolarization and a fast depolarizing input (for details see Methods, section Model assumption for the pons). The conductance of the cholinergic intrinsic currents regulates the degree of hyperpolarization in the resting potential of R-PBL neurons. As shown in Fig. S1, with a fixed strength of synaptic input from C-PBL neurons, only a carefully selected range of cholinergic conductance is able to trigger a calcium spike with bursts; too strong or weak resting hyperpolarization only cause a small depolarizing effect but no bursts. Therefore, based on such nonlinear properties, we are able to find an appropriate value of the cholinergic conductance to achieve the biologically-based temporal characteristics of pontine activity reported in Table S1.

### Validation of Ponto-thalamic projections

We consider one projection necessary if the PGO waveforms generated with it can not be reproduced in its absence, regardless of the variation of the other projections. In practice, this can be verified by using a dimensionality reduction technique mapping each shape to a point in a 2-dimensional space, and checking whether the waveforms simulated with and without each projection map to distinct regions of this space.

We thus complement the dataset of the original PGO waveforms generated with full ponto-thalamic projections (which underlies the Fig. S1 and Fig. 3) with 5 datasets of 500 substitute PGO waveforms by blocking each of the ponto-thalamic projections. Blockade is implemented by setting the corresponding projection strength to zero, while the strength of unblocked projections are set to randomly deviate maximally ±100% relative to the optimized values. We assume that the range of parameter variations is large enough to cover most of the parameter space for the generation of PGO waveforms. From these 6 datasets we take as features the waveforms of population membrane potential, firing rates and LFPs, separately for pre-REM and REM states. We reduce the high-dimensional feature space to 2 dimensions by applying T-SNE which has been shown to perform well on dimension reduction problems involving time series (Maaten and Hinton, 2008). The point clouds in Fig. S2 show the REM features of the optimized PGO waveforms with full projections are clearly separated from the 5 blockade conditions, while they are not during pre-REM states. This suggests that each of the ponto-thalamic projections has a specific role in the generation of PGO-related waveforms.

### Spindle event detection

Type-I spindles were detected by threshold crossings of the membrane potential of TC neurons at −45mV. The detected points above threshold were grouped into one event if the were within 2000ms away from each other, while events were aligned by the location of the maximum peak within the groups. Events were discarded if they oscillates greatly before and after the peak, which suggests as a mixture of type-I and type-II spindles. This is detected when the difference between maximum and minimum values between the membrane potential of TC neurons in two intervals ([−1500, −1000] and [1000, 1500]ms relative to the peak) is larger than 5mV.

Type-II spindles were selected by detection of the same thalamic signal between −55mV and −45mV. The alignment procedure is also the same as for type-I spindles. The mixture of type-I and type-II spindles are detected and discarded if the membrane of Pyr neurons stays below −65mV in a window of 6000ms around the peak (suggesting a K-complexes that corresponds to type-I spindles).

### Computation of triggered plasticity

The STDP weight variation defined in equation 1 were estimated for each PGO simulation sampled at 1000Hz as follows: (1) for all simulation times *t*, the convolution 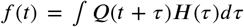, were estimated by discrete time convolution of the sampled STDP function truncated to a 600-ms time window centered around 0 by the sampled rate *Q_Pyr_*; (2) then the resulting time series were multiplied with the simulation of the presynaptic current *J_Syn–Syn_*(*t*), before smoothing it on a time window of 100ms to estimate the change of the synaptic strength 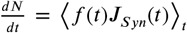. The peri-event changes of synaptic strength were aligned across 1000 PGO events to yield the statistics represented in Fig. 4D. The same procedure was applied for the case of spindles, combined with the spindle detection procedure described above.

### Python implementation

All simulation were implemented in the Python package ANNarchy^1^ (acronym for Artificial Neural Networks architect).

## Footnotes

1 https://www.tu-chemnitz.de/informatik/KI/projects/ANNarchy/index.php

## Notes

### Competing Interest Statement

The authors have declared no competing interest.

