## Supplementary Materials for "A model of Ponto-Geniculo-Occipital waves supports bidirectional control of cortical plasticity across sleep-stages"

**This PDF file includes:**

- Figures S1-S4
- Tables S1-S7
- Appendices A-C

### Supplementary Figures and Tables

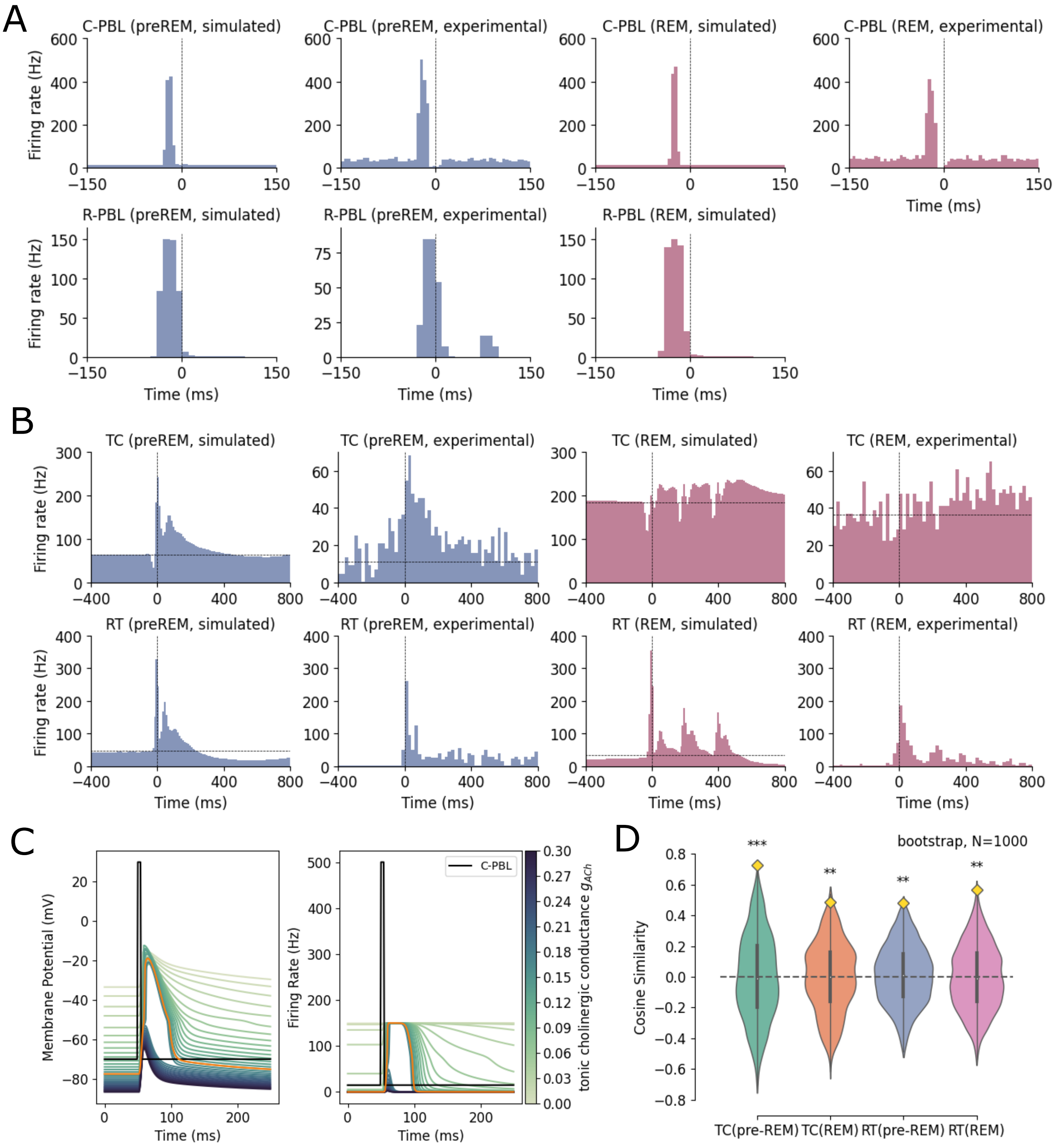


**Supplementary Figure S1.** Model Reproduction of Pontine and Thalamic neuronal activities. (A-B) Comparison of simulated and experimental pontine (A) and thalamic (B) peri-PGO histograms during pre-REM and REM states: in all conditions simulated results resemble experimental ones. The histograms are averaged over 1000 trials of simulated events with small variations of the ponto-thalamic projections. (C) ACh-tuned pontine neuronal activities. PGO-triggered membrane potential and firing rate of R-PBL neurons modulated by the conductance of a tonic cholinergic current. The conductance is critical to separate the neuronal activity into three patterns: with small conductance ($g_{ACh} <= 0.04$), the membrane is slightly depolarized; with moderate conductance ($g_{ACh} = [0.04, 0.18]$), a calcium spike rides on the PGO-triggered depolarization; when the conductance is set large ($g_{ACh} > 0.18$), the calcium spike disappears. This effect reflects the nonlinear intrinsic properties of the pontine T-current regulated by the cholinergic input current. The orange line marks the selected value of $g_{ACh} = 0.16$. (D) Cosine similarity as a measure of similarity between simulated and experimental PGO waves. Yellow diamonds represent the cosine similarity between the simulated and experimental peri-PGO histograms presented in (b); violin-plots show the bootstrapped distribution of cosine similarity calculated with 1000 epochs randomly-selected from the original simulation. Stars indicate that the cosine similarity in (b) is significantly different from the bootstrapped null distribution (***:p<0.001; **:p<0.01; *:p<0.05).

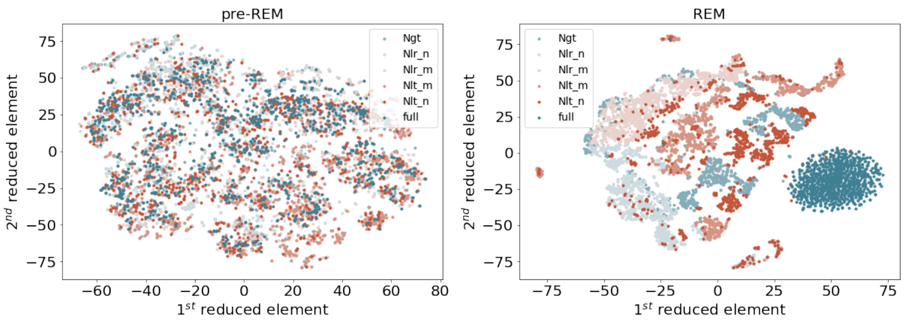


**Supplementary Figure S2.** Selective blockade of each ponto-thalamic projection alter PGO waveforms. The two subplots compare the separability of dimension-reduced features of optimal PGO waves (pink) and substitute ones obtained by blocking one ponto-thalamic projection at a time. (the blocked ones marked in the legend) in the simulated pre-REM (left panel) and REM stages (right panel). Substitute PGO waves are generated with large noise in the projection strength to cover the large parameter space. Notably, in REM stage, the isolated cluster of original PGO features show that the model is not able to generate the optimal PGO waveforms without any of the ponto-thalamic projections, suggesting the necessity of each projection.

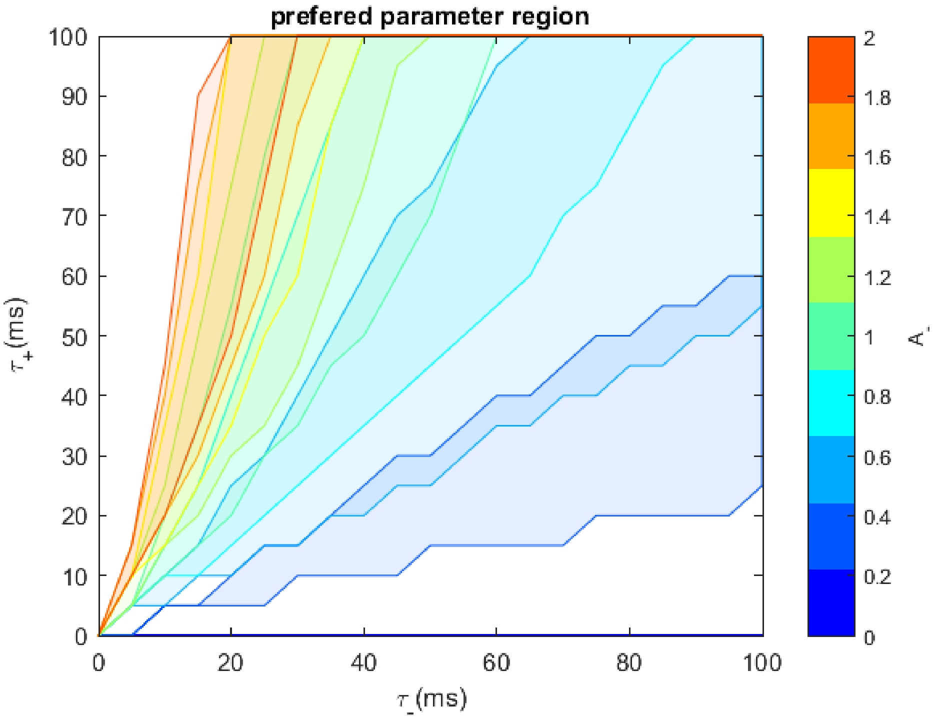


**Supplementary Figure S3.** Domain of the three STDP parameters for which the two PGO subtypes trigger opposite plasticity (LTP in pre-REM stage and LTD in REM stage). $A_-$ is scanned in the range of [0,2] with a step of 0.25. The time constants are still scanned in the range of [0,100]ms with a step of 5ms.

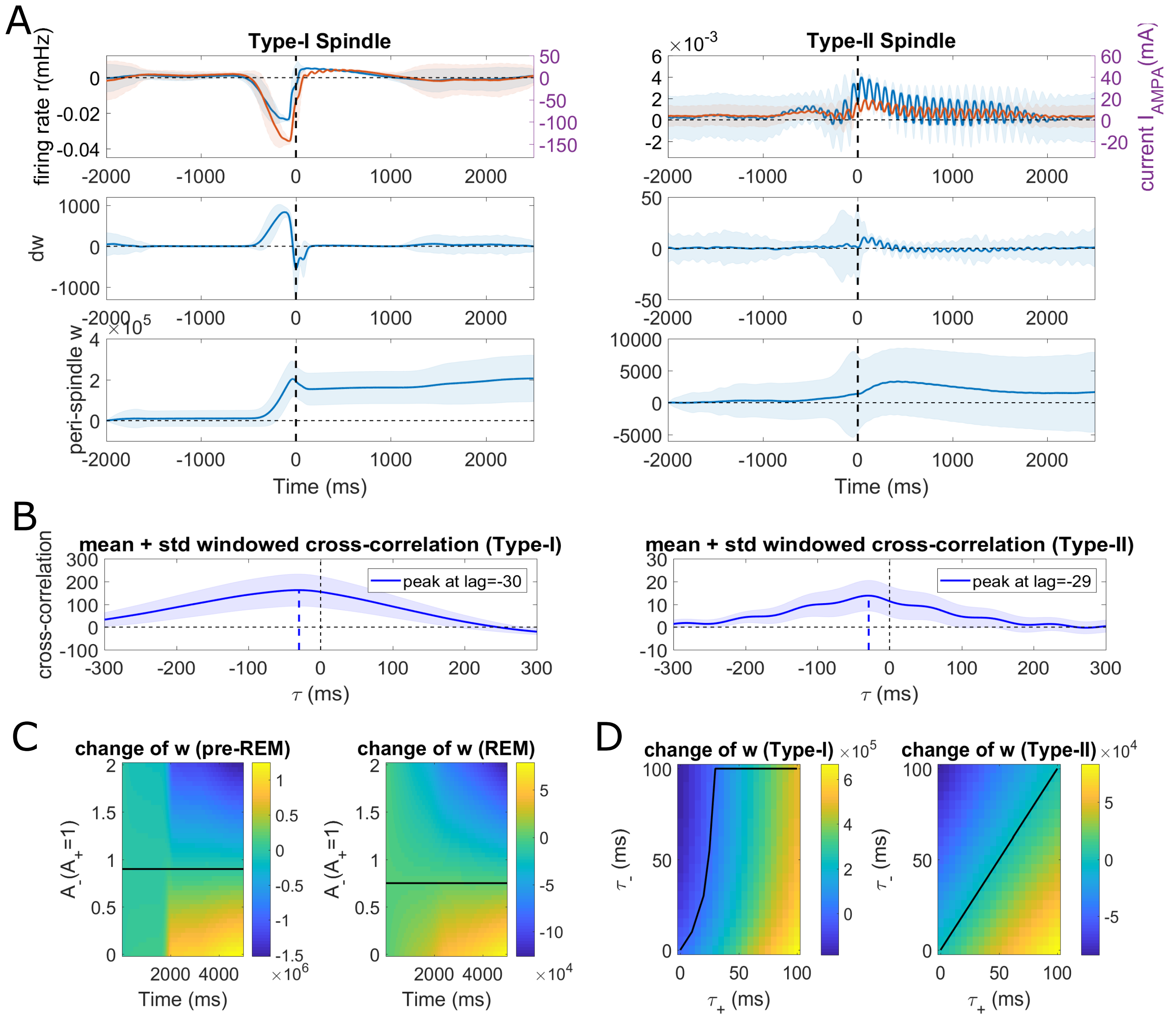


**Supplementary Figure S4.** Spindle-triggered cortical plasticity. (A) smoothed change of synatic strength induced by two subtypes of spindles. (top) demeaned waveforms of pre-synaptic current (red) and postsynaptic firing rate (blue). (middle) time-varying change of synaptic strength. (bottom) synaptic strength of the intra-cortical excitatory synapse changing with time. (B) Comparison of cross-correlation between the pre-synaptic current and the post synaptic activities. Blue shade represents standard deviation at each lag across events (N=1000). (C) Effect of STDP parameter $A^{-}$ on the plasticity direction induced by two spindle subtypes. Color bar indicates the relative increase/decrease of synaptic strength across time. Solid lines mark the critical value of the parameter that switches plasticity direction, i.e. LTP vs LTD; dashed lines correspond to the critical values of the other spindle subtype. (D) Effect of STDP parameter $\tau_+$ and $\tau_-$ on the plasticity direction induced by two PGO wave subtypes. Color bars and lines are analogous to C.

**Supplementary Table S1.** Quantitative similarities between pontine simulation and experimental results. See Datta and Hobson (1994) for characteristics related to C-PBL and Steriade et al. (1990) for that of R-PBL neurons.

| Electrophysioligical characteristic | simulated (ms) | experimental range (ms) |
|---|---|---|
| bursting duration (C-PBL) | 10 | [6, 16.7] |
| bursting duration (R-PBL) | 27 | [16.7, 41.7] |
| latency to bursting onset (C-PBL → RT) | 25 | [17.5, 32.5] |
| latency to bursting onset (R-PBL → RT) | 14 | [5, 15] |
| latency to negative peak in LFP (R-PBL → RT) | [35, 39] | [20, 40] |
| bursting interval (C-PBL) | set to 200 ms | around 200ms |

**Supplementary Table S2.** Parameter settings for neuron populations

| Symbol description | Symbol | Value | Unit |
|---|---|---|---|
| Commom parameters | | | |
| Membrane capacitance | $C_m$ | 1 | $\mu F/cm^2$ |
| Population of thalamocortical neurons ($t$) | | | |
| Maximal firing rate | $Q_t^{max}$ | $40 \cdot 10^{-3}$ | $ms^{-1}$ |
| Membrane time constant | $\tau_t$ | 20 | $ms$ |
| Firing threshold | $\theta_t$ | $-58.5$ | $mV$ |
| Default gain coefficient | $\sigma_t$ | 6 | $mV$ |
| Population of reticular thalamic neurons ($r$) | | | |
| Maximal firing rate | $Q_r^{max}$ | $400 \cdot 10^{-3}$ | $ms^{-1}$ |
| Membrane time constant | $\tau_r$ | 20 | $ms$ |
| Firing threshold | $\theta_r$ | $-58.5$ | $mV$ |
| Default gain coefficient | $\sigma_r$ | 6 | $mV$ |
| Population of intra-LG interneurons ($g$) | | | |
| Maximal firing rate | $Q_g^{max}$ | $170 \cdot 10^{-3}$ | $ms^{-1}$ |
| Membrane time constant | $\tau_c$ | 94 | $ms$ |
| Firing threshold | $\theta_c$ | $-58.5$ | $mV$ |
| Default gain coefficient | $\sigma_c$ | 4 | $mV$ |
| Population of transferring neurons ($l$) | | | |
| Maximal firing rate | $Q_r^{max}$ | $150 \cdot 10^{-3}$ | $ms^{-1}$ |
| Membrane time constant | $\tau_l$ | 20 | $ms$ |
| Firing threshold | $\theta_l$ | $-50.6$ | $mV$ |
| Default gain coefficient | $\sigma_l$ | 6 | $mV$ |
| Population of triggering neurons ($c$) | | | |
| Maximal firing rate | $Q_c^{max}$ | $500 \cdot 10^{-3}$ | $ms^{-1}$ |
| Membrane time constant | $\tau_c$ | 20 | $ms$ |
| Firing threshold | $\theta_c$ | $-58.5$ | $mV$ |
| Default gain coefficient | $\sigma_c$ | 6 | $mV$ |

**Supplementary Table S3.** Parameter settings for synaptic connections within thalamus

| Symbol description | Symbol | Value | Unit |
|---|---|---|---|
| Commom parameters | | | |
| Nernst potential of general leaky currents | $E_L$ | $-70.0$ | $mV$ |
| AMPA synapse in postsynaptic population $t$ $(at)$ | | | |
| Noise strength | $N_\phi$ | 0 | - |
| Synaptic rate constant | $\gamma_{at}$ | $100 \cdot 10^{-3}$ | $ms^{-1}$ |
| Nernst potential | $E_{at}$ | 0.0 | $mV$ |
| Noise variance | $\phi_n$ | 0.0 | - |
| GABA synapse in postsynaptic population $t$ $(gt)$ | | | |
| Connectivity constant | $N_{gt}$ | 5.0 | - |
| Synaptic rate constant | $\gamma_{gt}$ | $100 \cdot 10^{-3}$ | $ms^{-1}$ |
| Nernst potential | $E_{gr}$ | $-70.0$ | $mV$ |
| AMPA synapse in postsynaptic population $r$ $(ar)$ | | | |
| Connectivity constant | $N_{ar}$ | 3.0 | - |
| Synaptic rate constant | $\gamma_{ar}$ | $100 \cdot 10^{-3}$ | $ms^{-1}$ |
| Nernst potential | $E_{ar}$ | 0.0 | $mV$ |
| GABA synapse for self-feedback in population $r$ $(gr)$ | | | |
| Connectivity constant | $N_{gr}$ | 25.0 | - |
| Synaptic rate constant | $\gamma_{ar}$ | $100 \cdot 10^{-3}$ | $ms^{-1}$ |
| Nernst potential | $E_{ar}$ | $-70.0$ | $mV$ |

**Supplementary Table S4.** Parameter settings for synaptic connections from the Pons to thalamus

| Symbol description | Symbol | Value | Unit |
|---|---|---|---|
| nAChR synapse from population $l$ to population $t$ ($nt$) | | | |
| Connectivity constant | $N_{nt}$ | $0.1$ | - |
| Decay time | $\tau_{nt,1}$ | $27.8$ | $ms$ |
| Time constant | $\tau_{nt,2}$ | $1.69$ | $ms$ |
| Nernst potential | $E_{nt}$ | $-18.9$ | $mV$ |
| mAChR synapse from population $l$ to population $t$ ($mt$) | | | |
| Connectivity constant | $N_{mt}$ | $0.2$ | - |
| Synaptic rate constant | $\gamma_{mt}$ | $12.5 \cdot 10^{-3}$ | $ms^{-1}$ |
| Nernst potential | $E_{mt}$ | $-101$ | $mV$ |
| nAChR synapse from population $l$ to population $r$ ($nr$) | | | |
| Connectivity constant | $N_{nr}$ | $0.1$ | - |
| Decay time | $\tau_{nr,1}$ | $123.6$ | $ms$ |
| Time constant | $\tau_{nr,2}$ | $9.93$ | $ms$ |
| Nernst potential | $E_{nr}$ | $-5$ | $mV$ |
| mAChR synapse from population $l$ to population $r$ ($mr$) | | | |
| Connectivity constant | $N_{mr}$ | $0.4$ | - |
| Decay time | $\tau_{mr,1}$ | $639$ | $ms$ |
| Time constant | $\tau_{mr,2}$ | $92.09$ | $ms$ |
| Nernst potential | $E_{mr}$ | $-90$ | $mV$ |
| nAChR synapse from population $l$ to population $g$ ($ng$) | | | |
| Connectivity constant | $N_{ng}$ | $20$ | - |
| Synaptic rate constant | $\gamma_{ng}$ | $20 \cdot 10^{-3}$ | $ms^{-1}$ |
| Nernst potential | $E_{ng}$ | $-33$ | $mV$ |
| mAChR synapse from population $l$ to population $g$ ($mg$) | | | |
| Connectivity constant | $N_{mg}$ | $0.5$ | - |
| Decay time | $\tau_{mg,1}$ | $639$ | $ms$ |
| Time constant | $\tau_{mg,2}$ | $92.09$ | $ms$ |
| Nernst potential | $E_{mg}$ | $-90$ | $mV$ |
| AMPA synapse from population $c$ to population $l$ ($al$) | | | |
| Connectivity constant | $N_{al}$ | $2.95$ | - |
| Decay time | $\tau_{al,1}$ | $8.77$ | $ms$ |
| Time constant | $\tau_{al,2}$ | $1.66$ | $ms$ |
| Nernst potential | $E_{al}$ | $-3.4$ | $mV$ |
| NMDA synapse from population $c$ to population $l$ ($dl$) | | | |
| Connectivity constant | $N_{dl}$ | $0.44$ | - |
| Decay time | $\tau_{dl,1}$ | $129.4$ | $ms$ |
| Time constant | $\tau_{dl,2}$ | $2.02$ | $ms$ |
| Nernst potential | $E_{dl}$ | $16.3$ | $mV$ |

**Supplementary Table S5.** Parameter settings for synaptic connections

| Symbol description | Symbol | Value | Unit |
|---|---|---|---|
| Commom parameters | | | |
| Nernst potential of general leaky currents | $E_L$ | $-70.0$ | $mV$ |
| Nernst potential for T-currents | $E_{Ca}$ | $120.0$ | $mV$ |
| Nernst potential for $K^+$ currents | $E_K$ | $-100$ | $mV$ |
| $K^+$ leaky current in population $t$ | | | |
| Maximum conductance | $\overline{g}^t_{LK}$ | $0.041$ | $mS/cm^2$ |
| T-current in population $t$ | | | |
| Maximum conductance | $\overline{g}^t_T$ | $3.0$ | $mS/cm^2$ |
| h-current in population $t$ | | | |
| Maximum conductance | $\overline{g}_h$ | $0.051$ | $mS/cm^2$ |
| Conductivity scaling | $g_{inc}$ | $2$ | - |
| Calcium influx rate | $\alpha_{Ca}$ | $51.8 \cdot 10^{-6}$ | $mM/(mA \cdot ms)$ |
| Calcium time constant | $\tau_{Ca}$ | $10$ | $ms$ |
| Calcium resting state concentration | $Ca_0$ | $2.4 \cdot 10^{-4}$ | $ms^{-1}$ |
| Reaction velocity 1 of h-current | $k_1$ | $2.5 \cdot 10^7$ | $ms^{-1}$ |
| Reaction velocity 2 | $k_2$ | $4 \cdot 10^{-4}$ | $ms^{-1}$ |
| Reaction velocity 3 | $k_3$ | $1 \cdot 10^{-1}$ | $ms^{-1}$ |
| Reaction velocity 4 | $k_4$ | $1 \cdot 10^{-3}$ | $ms^{-1}$ |
| $K^+$ leaky current in population $r$ | | | |
| Maximum conductance | $\overline{g}^r_{LK}$ | $0.012$ | $mS/cm^2$ |
| T-current in population $r$ | | | |
| Maximum conductance | $\overline{g}^r_T$ | $2.3$ | $mS/cm^2$ |
| T-current in population $l$ | | | |
| Maximum conductance | $\overline{g}^l_T$ | $0.3$ | $mS/cm^2$ |
| Inward rectifier in population $l$ | | | |
| Maximum conductance | $\overline{g}^l_{IR}$ | $0.16$ | $mS/cm^2$ |

**Supplementary Table S6.** Parameter settings for connectivity strength in the generation of two types of PGO waves during pre-REM and REM

| Symbol description | Symbol | Value |
|---|---|---|
| isolated PGOs in pre-REM | | |
| nAChR current in TC neurons | $N_{nt}$ | $0.5$ |
| mAChR current in TC neurons | $N_{mt}$ | $0.15$ |
| nAChR current in RT neurons | $N_{nr}$ | $0.1$ |
| mAChR current in RT neurons | $N_{mr}$ | $0.5$ |
| isolated PGOs in REM and ACh-tuned PGOs in both states | | |
| nAChR current in TC neurons | $N_{nt}$ | $0.1$ |
| mAChR current in TC neurons | $N_{mt}$ | $0.05$ |
| nAChR current in RT neurons | $N_{nr}$ | $0.3$ |
| mAChR current in RT neurons | $N_{mr}$ | $0.4$ |

**Supplementary Table S7.** Parameter settings for $K^{+}$ leaky conductance in pre-REM and REM

| Symbol description | Symbol | Value | Unit |
|---|---|---|---|
| maximum $K^{+}$ leaky conductance during pre-REM | | | |
| for TC neurons | $\overline{g}^{t}_{LK,NREM}$ | 0.041 | $mS/cm^2$ |
| for TC neurons | $\overline{g}^{r}_{LK,NREM}$ | 0.012 | $mS/cm^2$ |
| maximum $K^{+}$ leaky conductance during pre-REM | | | |
| for TC neurons | $\overline{g}^{t}_{LK,REM}$ | 0.012 | $mS/cm^2$ |
| for TC neurons | $\overline{g}^{r}_{LK,REM}$ | 0.041 | $mS/cm^2$ |

### Appendix A: modeling assumptions

#### Origin of model choice

Our model focuses primarily on the reproduction of electrophysiological characteristics of PGO waves in the thalamus, where experimental PGO wave traces are most prominent and cellular mechanisms are relatively clear after extensive investigations in the field. Well-replicated thalamic PGO waves enable us to speculate on the effects PGO waves trigger in the cortex. We take as a starting point a neural mass model of thalamocortical network proposed in Schellenberger Costa et al. (2016), whose cortex and thalamus module, as well as the thalamic neuronal types (the TC and RT neurons), match well with our purpose. Then LGin neurons are added and connected with the pons in a biologically-realistic way.

This model is able to reproduce signatures of non-REM sleep, e.g. thalamic spindles and K-complexes in the cortex, by incorporating various intrinsic currents to generates specific neuronal activities (e.g. bursting). Thus, elaborating the model would also facilitate the investigation of interactions between PGO waves with these non-REM events.

To model the intrinsic properties of thalamic neurons, we follow the literature exploited in this thalamocortical model (Destexhe et al., 1996; Schellenberger Costa et al., 2016), i.e. a $K^+$ leaky current and a low-threshold Calcium T-current in both neurons populations, together with a hyperpolarization-activated anomalous rectifier h-current in TC neurons.

The cortex is simplified as a population of pyramidal cells interacting with a group of inhibitory neurons. Similarly, for the cortex, we keep the intrinsic current - a sodium-dependent potassium current $I_{KNa}$- in Pyr neurons so as to maintain the non-REM-related cortical oscillations (i.e. slow waves and K-complexes).

The addition of an LGin neurons follows a major hypothesis of thalamic PGO wave generation (Hu et al., 1989a). No extra intrinsic currents are associated to this population due to limited knowledge (Zhu et al., 1999).

#### Model assumption for the pons

Two groups of neurons in the pontine region, termed as the PGO *triggering neurons* and *transferring neurons*, are the executive elements of PGO waves (Datta, 1997). They are both located in the peribrachial area (PBL) (Silvestri and Kapp, 1998), which contains a number of important nuclei that are involved in regulation of sleep and arousal. Functionally, the PBL can be separated into two parts: the rostal (R-PBL) and the caudolateral (C-PBL) parts (Datta, 1997). The R-PBL mainly consists of the pedunculopontine tegmentum (PPT) nucleus and laterodorsal tegmentum (LDT) nucleus, while the most important nuclei in C-PBL include the parabrachial nucleus.

As their name indicates, experimental evidence showed that triggering neurons are PGO-state-on bursting neurons assumed to initiate the PGO phasic event located in C-PBL (Datta and Hobson, 1994; Datta, 1995, 1997). PGO triggering neurons were recorded to burst in high frequency (300-500Hz) $25 \pm 7$ ms before the thalamic PGO waves. These activities are hypothesized to propagate to the so-called transferring neurons in R-PBL, which are PGO-on low-frequency bursting neurons (Datta, 1997) firing low-frequency ($120\text{-}180Hz$) bursts with 3-5 spikes $5\text{-}15ms$ before the thalamic PGO waves (McCarley et al., 1978; Nelson et al., 1983; Steriade et al., 1990; Sakai and Jouvet, 1980). The transferring neurons were presumed to project directly to the thalamus (Sakai and Jouvet, 1980) as the last relay station of the local PGO-related circuits in the pons (Paré et al., 1990).

In the model, C-PBL activities (of the triggering neurons) are taken as a trigger that initializes the whole network activity. To match the high-frequency bursts pooled across a population, the firing rate of C-PBL neurons should rise rapidly and persist for a short period. It is then natural to model the C-PBL activity in pre-REM stage as a brief pulse lasting $10ms$ as an approximation of the bursts duration (Datta and Hobson, 1994). Deducing from experimentally-reported peri-PGO spike histograms (Datta and Hobson, 1994), we assume that REM-related C-PBL bursts can be modelled by three $10 - ms$ pulses with a bursting interval of $200ms$.

The R-PBL PGO-transferring neuron populations are modeled with the same type of sigmoidal activation function as for the thalamic neurons (Eq. 2). The activation threshold remain unchanged because the pontine and thalamic spikings are both associated with fast $Na^{2+}$ spikes and should match each other. We present briefly here the synaptic connection from C-PBL neurons to R-PBL neurons and some intrinsice cellular mechanisms of R-PBL neurons.

The projection from triggering neurons to transferring neurons is presumably glutamatergic (Inglis and Semba, 1996; Sanchez and Leonard, 1994, 1996; Datta, 2012), indirectly supported by the findings that other neurotransmitters are inhibitory (Luebke et al., 1992; Leonard and Llinás, 1994; Williams and Reiner, 1993). Electrophysiological studies showed the projection could be mediated by a combination of NMDA and AMPA receptors (Sanchez and Leonard, 1996), with a contribution ratio as NMDA:AMPA = 1:5. The same experiment also quantified the decay times for both currents: 8.77ms for the AMPA channel and 129.4ms for for NMDA. The Nernst potentials were measured to be 16.3mV for NMDA and 3.4mV for AMPA (close to the theoretical value 0mV).

Conductance of the AMPA channel is invariant to the postsynaptic membrane potential. Non-linearity in NMDA currents has long been reported and well-documented. We followed the classical modelling of the voltage dependence first introduced by Destexhe ((Destexhe et al., 1994)):

$$g_{NMDA}(V_l) = \frac{1}{1 + \exp(-0.0062V_l)}[Mg^{2+}]_o/3.57 \tag{26}$$

where $l$ represents the population of R-PBL transferring neurons, and $V_l$ denotes its membrane potential. $[Mg]_o$ represents the extracellular concentration for $Mg^{2+}$ ions.

The slow-frequency bursts occuring in transferring neurons are rebound bursts evoked by activating an intrinsic low-threshold calcium T-current under hyperpolarization (Kang and Kitai, 1990; Kamondi et al., 1992). As it is similar to the thalamic T-currents discovered in the TC neurons, we modelled it with Eq. 5 (Destexhe et al., 1998, 1994), but refitted the conductance with digitized experimental I-V curves (Kamondi et al., 1992). We constructed a Boltzmann-like function that is able to approximate the activation curve:

$$m^l_\infty(V_l) = \frac{2}{1 + \exp(-(V_l + 50.6/0.44) + \exp((V_l + 50.6/17.4}} \tag{27}$$

In a similar way, we fitted the inactivation gating variable leading to:

$$h^l_\infty(V_l) = \frac{1}{1 + \exp((V_l + 65)/2.7)} \tag{28}$$

Considering the increasing concentration of ACh during PGO-related sleep stages, together with evidence of the inhibitory effects of Ach on transferring neurons (Leonard and Llinás, 1994), we assume that some cholinergic modulatory inputs causes the hyperpolarization as a prerequisite to de-inactivate the T-current. The cholinergic input goes through a potassium inward rectifier mediated by a muscarinic receptor (Leonard and Llinás, 1994). The I-V curve of cholinergic influence has already been characterized, from which we defined an approximate mathematical formulation with a procedure similar to the fitting of T-currents.

$$g_{IR}(V_l) = \frac{1}{1 + \exp((V_l + 35)/10.9)} \tag{29}$$

#### Propagation of pontine PGO waves to the thalamus

The complete model involving carefully-designed ponto-thalamic synaptic connections is illustrated in Fig. 1B. After receiving bursting activities from the triggering neurons, the transferring neurons send cholinergic inputs to the three thalamic neurons (TC and RT neurons and LGin neurons) via 5 cholinergic ponto-thalamic projections. Here we briefly present our assumptions regarding the chemical nature and kinetics of the pontine-thalamic projections and neurophysiological evidence supporting them.

The R-PBL neurons send two excitatory projections to the TC neurons via cholinergic projections mediated by both nAChR and mAChR receptors. The nAChR-based channel, underlying the generation mechanism of a fast depolarization in deafferented cats (McCormick and Prince, 1987b; Hu et al., 1988, 1989a,a) and short bursting in naturally sleeping cats (Steriade et al., 1989; McCormick and Prince, 1987b; Hu et al., 1989b), let through a mixed cation current with a voltage-independent conductance and a Nernst potential of $18.9 \pm 8.9mV$ (McCormick and Prince, 1987b). The mAChR-mediated synapse, generating the prolonged spiking in TC neurons following the initial bursts, is coupled to a leaky $K^+$ channel via G-protein cascade (reversal potential: $-97 \pm 6.1mV$, note that the term "leaky" entails a voltage-independent conductance) (McCormick, 1992). Specifically, we speculate that this potassium leaky channel is the same channel as the $K^+$ leaky channel already included in the original model of TC neurons (see Section Intrinsic currents and Section Cholinergic modulation of PGO waves), whose role is designed to mediate membrane de-/hyperpolarization (McCormick and Prince, 1987b; Bista et al., 2012, 2015). We introduced a saturation mechanism: the maximum amount of conductance decrease caused by the mAChR-mediated synapse is equal to its resting conductance, i.e. it cannot go beyond complete closure. This saturation mechanism was not reported but implied in the experimental papers, and has been proven useful in replicating the switch between PGO wave subtypes (see Fig. S1).

Clear evidence suggests that the RT neurons also receive pontine inputs via both nAChR- and mAChR-mediated synapses, corresponding to a PGO-triggered fast depolarization/bursting and slow hyperpolarization observed in RT neurons (Hu et al., 1989a; Lee and McCormick, 1995; Oda et al., 2007; Sun et al., 2013; Beierlein, 2014). The time constants of synatic kinetics were quantified (rise time: $10.8ms$, decay time: $123.6ms$), enabling us to apply the 'two-exponential' form of synaptic model. Following classical models of nAChR-mediated channels, we assume a linear voltage-independent with a reversal potential of $-5mV$. On the contrary, the mAChR-regulated channel, associated with a $K^+$ channel with the Nernst potential of around $-93.2 \pm 0.6mV$ (Sun et al., 2013), works as a inward rectifier (Hu et al., 1989a; Oda et al., 2007; Lee and McCormick, 1995; Sun et al., 2013; Beierlein, 2014), whose conductance decreases with increased membrane potential, with voltage dependence characterized and fitted with asigmoidal function:

$$g_{mAChR}(V_r) = \frac{1}{1 + \exp((V_r + 66.3)/29.1)} \tag{30}$$

The corresponding rise time and decay time of the synaptic kinetics are $107.6 \pm 8.6ms$ and $639.0 \pm 102ms$.

The assumption that LGin neurons contribute to thalamic PGO wave generation is supported by the existence of a transient hyperpolarization of TC neurons caused by depolarization in LGin neurons (Hu et al., 1989a,b,c; Steriade et al., 1989). We model a nAChR-mediated projection from R-PBL neurons to LGin neurons to generate the depolarization (Zhu et al., 1999), with the same model as the corresponding channel in TC neurons with differently-tuned synaptic kinetics. Conservatively, for simplicity we omit the other potentially existing instrinsic current, as current experimental evidence is insufficient to support their roles in thalamic PGO wave formation.

The transmission of PGO-related activities from pons to the thalamus is not instataneous but delayed by membrane and axonal properties. For the 5 pontine-thalamic connections, the only precisely reported fact is the latency difference between nAChR-mediated and mAChR-mediated currents in RT neurons, as $28ms$ (Sun et al., 2013). Latency histograms of the nAChR-mediated currents in TC and RT neurons (Hu et al., 1989c) also suggest a constraint in setting the delays. Extrapolating with all taken into consideration, we implement the delays in our model by setting a fixed latency for each projection that is consistent with the experimentally-revealed facts.

#### Cholinergic modulation of PGO waves

The transition from pre-REM to REM sleep stages is strongly dependent on the changes of certain neuromodulators (Datta, 1997). To reproduce the difference between PGO wave subtypes during pre-REM and REM, we need to know how neuromodulation affects the PGO propagating network.

Modulatory neuron populations associated with ACh and monoamines (serotonin (5-HT) and noradrenaline (NE)) are reciprocally interacting to influence the activities of the several neuron types related to PGO wave generation (Hobson, 2009). Wakefulness and non-REM sleep is accompanied with high concentration of monoamines and low concentration of ACh. In the transitional stage (i.e. pre-REM), the activities of aminergic neurons decrease while the cholinergic neurons are gradually activated. When REM sleep is approached, cholinergic activities remain persistently in high level whereas aminergic activities are suppressed. In short, aminergic neurons plays a disinhibitory gating role for the cholinergic neurons, i.e. activities of the former is negatively correlated with the latter. Thus for simplicity we can omit the monoamines and model only the effect of ACh on the network activities.

### Appendix B: Derivation of synapse representation

#### Alpha function

For a synapse current (type $m$) from presynaptic population $k'$ to postsynaptic population $k$ with connectivity strength $N_{k'k}$, assume its impulse response is an alpha function:

$$h_m(t) = \gamma_m^2 \cdot t \cdot \exp(-\gamma_m t) \tag{31}$$

If we perform a Laplace transform to the impulse response, we would obtain:

$$\mathcal{L}[h_m(t)](s) = \mathcal{L}[\gamma_m^2 t \exp(-\gamma_m t)](s) = \gamma_m^2 \cdot \frac{\Gamma(2)}{(s+\gamma_m)^2} = \frac{\gamma^2}{(s+\gamma_m)^2} \tag{32}$$

Here the impulse response equals:

$$h_m(t) = \frac{s_{mk}(s)}{N_{k'k} \cdot Q_{k'}(s)} = \frac{\gamma_m^2}{s^2 + 2\gamma_m s + \gamma_m^2} \tag{33}$$

If we rearrange the terms, we can get:

$$s^2 \cdot s_{mk}(s) + 2s \cdot s_{mk}(s) + \gamma_m^2 \cdot s_{mk}(s) = \gamma_m^2 \cdot N_{k'k} \cdot Q_{k'}(s) \tag{34}$$

After an inverse Laplace transform and a rearrangement of the terms, we can reach the differential equation that we want:

$$\ddot{s}_{mk} = \gamma_m^2 (N_{k'k} \cdot Q_{k'}(V_{k'}(t)) - s_{mk}) - 2\gamma_m \dot{s}_{mk} \tag{35}$$

#### 'Two-exponential' inpulse response function

Similarly, for the 'two-exponential' type of synaptic impulse response:

$$h_m(t) = B(\exp(-t/\tau_1) - \exp(-t/\tau_2)) \tag{36}$$

The Laplace transformed $h_m(t)$ would take the following form:

$$\mathcal{L}[h_m(t)](s) = \mathcal{L}[B(\exp(-t/\tau_1) - \exp(-t/\tau_2))](s) = \frac{B}{s+\tau_1^{-1}} - \frac{B}{s+\tau_2^{-1}} \tag{37}$$

Then we include the input and output of the system in the Laplace domain:

$$\frac{s_{mk}(s)}{N_{k'k} \cdot Q_{k'}(s)} = \frac{B(\tau_2^{-1} - \tau_1^{-1})}{s^2 + (\tau_2^{-1} + \tau_1^{-1})s + \tau_2^{-1}\tau_1^{-1}} \tag{38}$$

If we rearrange the terms, we can get:

$$s^2 \cdot s_{mk}(s) + (\tau_2^{-1} + \tau_1^{-1})s \cdot s_{mk}(s) + \tau_2^{-1}\tau_1^{-1} \cdot s_{mk}(s) = B(\tau_2^{-1} - \tau_1^{-1}) \cdot N_{k'k} \cdot Q_{k'}(s) \tag{39}$$

After an inverse Laplace transform and a rearrangement of the terms, The differential version is obtained after a reverse Laplace transform

$$\ddot{s}_{mk} = B(\tau_2^{-1} - \tau_1^{-1}) N_{k'k} \cdot Q_{k'}(V_{k'}(t)) - \tau_2^{-1}\tau_1^{-1} s_{mk} - (\tau_2^{-1} + \tau_1^{-1})\dot{s}_{mk} \tag{40}$$

### Appendix C: Full list of equations

#### Neuron populations

Firing rate function of population $k$, where $k \in \{t, r, g, l, c\}$, representing the TC neurons, the RT neuron, the intra-LG interneurons, the PGO transferring neurons and the PGO triggering neurons.

$$Q_k = \frac{Q_k^{max}}{1 + \exp(-(V_k - \theta_k)/\sigma_k)} \tag{41}$$

Membrane potential adaptations for population $t, r, g, l$

$$\begin{aligned}\tau_t \dot{V}_t = \quad & -J_L^t - J_{AMPA}(s_{at}) - J_{nAChR}(s_{nt}) - J_{GABA}(s_{gt}) - \\ & C_m^{-1} \tau_t (I_{LK}^t + I_T^t + I_h)\end{aligned} \tag{42}$$

$$\tau_r \dot{V}_r = -J_L^r - J_{AMPA}(s_{ar}) - J_{GABA}(s_{gr}) - J_{nAChR}(s_{nr}) - J_{mAChR}(s_{mr}) - C_m^{-1} \tau_r (I_{LK}^r + I_T^r) \tag{43}$$

$$\tau_g \dot{V}_g = -J_L^g - J_{nAChR}(s_{ng}) - J_{mAChR}(s_{ng}) \tag{44}$$

$$\tau_l \dot{V}_l = -J_L^l - J_{AMPA}(s_{al}) - J_{NMDA}(s_{dl}) - C_m^{-1} \tau_l (I_T^l + I_{ACh}) \tag{45}$$

#### Synaptic currents

Synaptic currents are denoted in the form of $J(s_{ij})$. $i \in \{a, g, n, m, d\}$ states the synaptic type AMPA, GABA, nAChR, mAChR and NMDA, whereas $j \in \{t, r, g, l, c\}$ indicates the postsynaptic population in Section .

General leaky currents of population $k$, where $k \in \{t, r, g, l, c\}$

$$J_L^k = (V_k - E_L^k) \tag{46}$$

AMPA synapse in postsynaptic population $t$

$$J_{AMPA}(s_{at}) = s_{at} \cdot (V_r - E_{at}) \tag{47}$$

$$\ddot{s}_{at} = \gamma_{at}^2 (\phi_n - s_{at}) - 2\gamma_{at} \dot{s}_{at} \tag{48}$$

GABA synapse in postsynaptic population $t$

$$J_{GABA}(s_{gt}) = s_{gt} \cdot (V_t - E_{gt}) \tag{49}$$

$$\ddot{s}_{gt} = \gamma_{gt}^2 \cdot \left( N_{gt} \cdot Q_r(V_r) + N_{gt} \cdot Q_g(V_g) - s_{gt} \right) - 2\gamma_{gt} \dot{s}_{gt} \tag{50}$$

nAChR synapse in postsynaptic population $t$

$$J_{nAChR}(s_{nt}) = s_{nt} \cdot (V_t - E_{nt}) \tag{51}$$

$$\ddot{s}_{nt} = B_{nt} (\tau_{nt,2}^{-1} - \tau_{nt,1}^{-1}) N_{nt} Q_l(V_l) - \tau_{nt,2}^{-1} \tau_{nt,1}^{-1} s_{nt} - (\tau_{nt,2}^{-1} + \tau_{nt,1}^{-1}) \dot{s}_{nt} \tag{52}$$

mAChR synapse in postsynaptic population $t$

$$J_{mAChR}(s_{mt}) = s_{mt} \cdot (V_t - E_{mt}) \tag{53}$$

$$\ddot{s}_{mt} = \gamma_{mt}^2 \left( N_{mt} \cdot Q_l(V_l) - s_{mt} \right) - 2\gamma_{mt} \dot{s}_{mt} \tag{54}$$

AMPA synapse in postsynaptic population $r$

$$J_{AMPA}(s_{ar}) = s_{ar} \cdot (V_r - E_{ar}) \tag{55}$$

$$\ddot{s}_{ar} = \gamma_{ar}^2 \left( (N_{ar}) \cdot Q_t(V_t) - s_{ar} \right) - 2\gamma_{ar}\dot{s}_{ar} \tag{56}$$

GABA synapse for self-feedback in population $r$

$$J_{GABA}(s_{gr}) = s_{gr} \cdot (V_r - E_{gr}) \tag{57}$$

$$\ddot{s}_{gr} = \gamma_{gr}^2 \left( (N_{gr}) \cdot Q_r(V_r) - s_{gr} \right) - 2\gamma_{gr}\dot{s}_{gr} \tag{58}$$

nAChR synapse in postsynaptic population $r$

$$J_{nAChR}(s_{nr}) = s_{nr} \cdot (V_r - E_{nr}) \tag{59}$$

$$\ddot{s}_{nr} = B_{nr}(\tau_{nr,2}^{-1} - \tau_{nr,1}^{-1}) N_{nr} Q_l(V_l) - \tau_{nr,2}^{-1}\tau_{nr,1}^{-1} s_{nr} - (\tau_{nr,2}^{-1} + \tau_{nr,1}^{-1})\dot{s}_{nr} \tag{60}$$

mAChR synapse in postsynaptic population $r$

$$J_{mAChR}(s_{mr}) = s_{mr} \cdot g_{mAChR}(V_r) \cdot (V_r - E_{mr}) \tag{61}$$

$$g_{mAChR}(V_r) = \frac{1}{1 + \exp((V_r + 66.3)/29.1)} \tag{62}$$

$$\ddot{s}_{mr} = B_{mr}(\tau_{mr,2}^{-1} - \tau_{mr,1}^{-1}) N_{mr} Q_l(V_l) - \tau_{mr,2}^{-1}\tau_{mr,1}^{-1} s_{mr} - (\tau_{mr,2}^{-1} + \tau_{mr,1}^{-1})\dot{s}_{mr} \tag{63}$$

nAChR synapse in postsynaptic population $g$

$$J_{nAChR}(s_{ng}) = s_{ng} \cdot (V_g - E_{ng}) \tag{64}$$

$$\ddot{s}_{ng} = B_{ng}(\tau_{ng,2}^{-1} - \tau_{ng,1}^{-1}) N_{ng} Q_l(V_l) - \tau_{ng,2}^{-1}\tau_{ng,1}^{-1} s_{ng} - (\tau_{ng,2}^{-1} + \tau_{ng,1}^{-1})\dot{s}_{ng} \tag{65}$$

mAChR synapse in postsynaptic population $g$

$$J_{mAChR}(s_{mg}) = s_{mg} \cdot (V_g - E_{mg}) \tag{66}$$

$$\ddot{s}_{mg} = B_{mg}(\tau_{mg,2}^{-1} - \tau_{mg,1}^{-1}) N_{mg} Q_l(V_l) - \tau_{mg,2}^{-1}\tau_{mg,1}^{-1} s_{mg} - (\tau_{mg,2}^{-1} + \tau_{mg,1}^{-1})\dot{s}_{mg} \tag{67}$$

AMPA synapse in postsynaptic population $l$

$$J_{AMPA}(s_{al}) = s_{al} \cdot (V_l - E_{al}) \tag{68}$$

$$\ddot{s}_{al} = B_{al}(\tau_{al,2}^{-1} - \tau_{al,1}^{-1}) N_{al} Q_c(V_c) - \tau_{al,2}^{-1}\tau_{al,1}^{-1} s_{al} - (\tau_{al,2}^{-1} + \tau_{al,1}^{-1})\dot{s}_{al} \tag{69}$$

NMDA synapse in postsynaptic population $l$

$$J_{NMDA}(s_{dl}) = s_{dl} \cdot g_{NMDA}(V_l)(V_r - E_{dl}) \tag{70}$$

$$g_{NMDA}(V_l) = \frac{1}{1 + \exp(-0.0062 V_l)} [Mg^{2+}]_o / 3.57 \tag{71}$$

$$\ddot{s}_{dl} = B_{dl}(\tau_{dl,2}^{-1} - \tau_{dl,1}^{-1}) N_{dl} Q_c(V_c) - \tau_{dl,2}^{-1}\tau_{dl,1}^{-1} s_{dl} - (\tau_{dl,2}^{-1} + \tau_{dl,1}^{-1})\dot{s}_{dl} \tag{72}$$

#### Intrinsic currents

Intrinsic currents are presented as $I_i^k$, with the subscription $i \in \{LK, T, h, IR\}$ describing the current types ($K^+$ leaky currents, T-currents, h-currents and $k^+$ inward rectification currents). The superscription indicates the neuron population where the intrinsic currents are located.

Potassium leaky currents of population $t$

$$I_{LK}^t = \overline{g}_{LK}^t \cdot (V_t - E_K) \tag{73}$$

Low threshold Calcium T-current for population $t$

$$I_T^t = \overline{g}_T^t \cdot (m_\infty^t)^2 \cdot h_T^t \cdot (V_t - E_{Ca}) \tag{74}$$

$$m_\infty^t = \frac{1}{1 + \exp(-(V_t + 59)/6.2)} \tag{75}$$

$$\dot{h}_T^t = (h_\infty^t - h_T^t)/\tau_h^t \tag{76}$$

$$h_\infty^t = \frac{1}{1 + \exp(-(V_t + 81)/4)} \tag{77}$$

$$\tau_h^t = (30.8 + (211.4 + \exp((V_t + 115.2)/5))/(1 + \exp((V_t + 86)/3.2)))/3^{1.2} \tag{78}$$

Anomalous inward rectifier h-current for population $t$

$$I_h^t = \overline{g}_h \cdot (m_{h1} + g_{inc} m_{h2}) \cdot (V_t - E_h)) \tag{79}$$

$$\dot{m}_{h1} = (m_\infty^h (1 - m_{h2}) - m_{h1})/\tau_m^h - k_3 * P_h m_{h1} + k_4 m_{h2} \tag{80}$$

$$\dot{m}_{h2} = k_3 P_h m_{h1} - k_4 m_{h2} \tag{81}$$

$$P_h = k_1 [Ca]^4/(k_1 [Ca]^4 + k_2) \tag{82}$$

$$[\dot{Ca}] = -\alpha_{Ca} I_T^t - ([Ca] - Ca_0)/\tau_{Ca} \tag{83}$$

Potassium leaky currents of population $r$

$$I_{LK}^r = \overline{g}_{LK}^r \cdot (V_r - E_K) \tag{84}$$

Low threshold Calcium T-current for population $r$

$$I_T^r = \overline{g}_T^r \cdot \left(m_\infty^r\right)^2 \cdot h_T^r \cdot (V_r - E_{Ca}) \tag{85}$$

$$m_\infty^r = \frac{1}{1 + \exp(-(V_r + 52)/7.4)} \tag{86}$$

$$\dot{h}_T^r = (h_\infty^r - h_T^r)/\tau_h^r \tag{87}$$

$$h_\infty^r = \frac{1}{1 + \exp(-(V_r + 80)/5)} \tag{88}$$

$$\tau_h^t = (85 + 1/(\exp((V_r + 48)/4))/(1 + \exp(-(V_r + 407)/50)))/3^{1.2} \tag{89}$$

Low threshold Calcium T-current for population $l$

$$I_T^l = \overline{g}_T^l \cdot \left(m_\infty^l\right)^2 \cdot h_T^l \cdot (V_l - E_{Ca}) \tag{90}$$

$$m^l_\infty = \frac{1}{1 + \exp(-(V_l + 50.6)/0.44) + \exp((V_l + 50.6)/17.4)} \tag{91}$$

$$\dot{h}^l_T = (h^l_\infty - h^l_T)/\tau^l_h \tag{92}$$

$$h^r_\infty = \frac{1}{1 + \exp((V_l + 65)/2.7)} \tag{93}$$

$$\tau^t_h = (52 + (211.4 + \exp((V_l + 115.2)/5))/(1 + \exp((V_l + 86)/3.2)))/\exp(1.2\log(3)) \tag{94}$$

$K^+$ Inward rectifier of cholinergic input in population $l$

$$I^l_{IR} = \overline{g}^l_{IR} \cdot g_{IR}(V_l) \cdot (V_l - E_K) \tag{95}$$

$$g^l_{IR} = \frac{1}{1 + \exp((V_l + 66.3)/29.1)} \tag{96}$$

#### State modulation

$$\Delta[ACh](t) = -\cos(\frac{2\pi}{T}t) \tag{97}$$

$$\overline{g}^t_{LK}(t) = -0.5(\overline{g}^t_{LK,NREM} - \overline{g}^t_{LK,REM}) \cdot (\Delta[ACh](t) - 1) + \overline{g}^t_{LK,REM}) \tag{98}$$

$$\overline{g}^r_{LK}(t) = 0.5(\overline{g}^r_{LK,REM} - \overline{g}^r_{LK,NREM}) \cdot (\Delta[ACh](t) + 1) + \overline{g}^r_{LK,NREM}) \tag{99}$$
